# CebraEM: A practical workflow to segment cellular organelles in volume SEM datasets using a transferable CNN-based membrane prediction

**DOI:** 10.1101/2023.04.06.535829

**Authors:** Julian Hennies, José Miguel Serra Lleti, Constantin Pape, Sultan Bekbayev, Viktoriia Gross, Anna Kreshuk, Yannick Schwab

## Abstract

Segmentation of large-volume datasets obtained by volume SEM techniques is a challenging task that generally requires a considerable amount of human effort. Despite recent advances in deep learning leading to the successful segmentation of cellular organelles in a variety of datasets, it is still challenging and time-consuming to produce the necessary data for training a convolutional neural network as well as to set up targeted post-processing pipelines to obtain a good quality full-volume semantic instance segmentation. We present CebraEM, a software package that uses a novel workflow for the segmentation of organelles in volume EM datasets, which helps to minimize the annotation time for the generation of training data. It relies on a generic CNN-based membrane prediction, followed by a well-established machine-learning pipeline that includes over-segmentation before random forest classification and graph multi-cut grouping. The workflow was tested for the segmentation of organelles on different datasets originating from various sample preparations and imaging modalities in volume SEM, in each case resulting in state-of-the-art semantic instance segmentations without additional post-processing. Importantly, by considerably simplifying the segmentation problem, CebraEM empowers single users with the ability to efficiently segment hundreds of gigabytes of data.

## Introduction

In recent years, image data production in the biological sciences has witnessed dramatic growth. In Electron Microscopy (EM), the necessity of acquiring large volumes of tissue with the highest possible detail ignited the interest in volume-based EM methods acquired by Scanning Electron Microscopy (SEM) (1–3). In comparison to volume datasets obtained by transmission EM tomography-based approaches, volume SEM datasets achieve lower resolution yet a substantially larger field of view. The most popular volume SEM techniques are array tomography (4, 5), SBF-SEM (6), and FIB-SEM (7, 8). FIB-SEM can offer up to 4 nm iso-voxel resolution covering entire cells (9), while SBF-SEM can cover a large field of view, up to entire multicellular organisms (10) although compromising on slice thickness. In general, volume SEM is used to capture a complete anatomic landscape at the nanometer scale, making it a suitable tool to study organelle morphology and organelle-organelle interactions. Thus, it can facilitate a better understanding of cellular processes, such as the dynamics of the endoplasmic reticulum (ER) (11), its contact sites to other organelles (with membrane distances of 20 to 100 nm) (12, 13), or viral replication factories driving viral production in infected cells (14).

After acquisition and image pre-processing, segmentation is the most common procedure to lay the foundation for quantitative morphometric analysis. By segmentation, we understand the grouping of an individual set of voxels into separated regions, following object boundaries within the dataset. In semantic segmentation, clusters of pixels are assigned to a specific label, e.g., the organelle to be segmented. If in addition to tagging pixels into labels, there is a differentiation between individual objects (instances) of the same category in the current scene, we then talk about semantic instance segmentation.

Segmentation for specific organelles, such as the ER or mitochondria, allows volumetric quantification and visualization of separated objects into rendered projections. In EM, region boundaries appear naturally with cell membranes, which are exceptionally enhanced as the result of optimized sample preparations (2). With the growth of data over the recent years, and even though semi-automated solutions have been developed, segmentation has become tedious and highly time-consuming. Before the popularization of machine learning, only manual (15) or semi-automatic approaches (16, 17) aspired to segment complex organelles.

Segmentation in biology and medicine followed the historical trend of computer vision, with algorithms growing progressively in complexity towards machine and deep learning (DL) (18, 19). In EM, they were substantially driven by the neurobiology community and their need for improving tools for connectomics (20–22). In cell biology, the attempts at fully automated organelle segmentation were mostly enabled by convolutional neural networks (CNNs) like the 3D U-net (23). In most cases, the efforts focused on obtaining a high segmentation quality for just one organelle, like mitochondria (24–27), or nuclear membrane (17). More recently, DL methods were successfully applied to full organelle segmentation (28, 29).

Using DL for segmenting organelles in EM comes with a set of additional challenges. First, it requires an expert to generate a large corpus of annotated data, which will be used as ground truth during the training of the neural network. The network training process is often not very stable with a high risk of overfitting. Additionally, models for specific tasks usually do not generalize to new datasets (e.g., due to variation in sample preparation, resolution, contrast/brightness). Since increasing the training and labeled data to have better coverage of the target domain is sometimes difficult or not possible, other strategies have been developed.

In order to train models on the target domain, targeted or efficient ground truth annotation approaches are employed. For example, model-assisted labeling or active learning (30), uses a strategy of repeatedly annotating, training, and predicting a supervised model to iteratively converge to satisfactory results. Within each iteration, the output is curated manually to generate a larger ground truth set for the next training step.

If the dataset is challenging, this iteration process may prolong indefinitely, making it as arduous a task as manual labeling. Another way to increase the efficiency of ground truth annotation is the usage of semi-automated approaches (13, 31). To further facilitate the training procedure and simplify setting up deep learning models, many approaches introduce pretrained network architectures (28, 29) or create readily implemented frameworks for training (32–35). Despite those efforts, training deep learning models effectively still requires a certain level of knowledge in machine learning, which might discourage non-experts from training with their own data. Moreover, whereas organelle prediction maps generated by deep-learning-based approaches can yield high accuracies and enable the segmentation of a large variety of intra-cellular structures, none of the most recent organelle segmentation approaches circumvent the need for adapted post-processing efforts.

With CebraEM, we introduce a machine-learning-based workflow, derived from the multicut pipeline (20, 36) that tackles the challenges mentioned above. CebraEM circumvents the need for training DL models entirely and generates automated semantic instance segmentations without the requirement for post-processing steps. The foundation of the CebraEM workflow is to split the segmentation task into first the prediction of organelle boundaries and then the detection of the target organelles. The prediction of boundaries is performed with a pretrained CNN model (CebraNET) which, without retraining, predicts organelle boundaries and enables the partitioning of the data into supervoxels. The partitioning of the data reduces the complexity of the overall problem, enabling the usage of a shallow model as well as substantially increasing the speed of ground truth annotation. For the ground truth annotation of selected regions at the super-voxel level, we implemented CebraANN, a plugin for napari (37).

Once annotated and trained, the CebraEM inference workflow can generate a suitable semantic instance segmentation of the full cell within a few hours per organelle.

To prove that our method can be generalized, we tested this segmentation workflow with several publicly available datasets covering a significant diversity in sample preparations method and volume SEM imaging modalities (see Methods, section Datasets): Firstly, we used a HeLa cell prepared by high-pressure freezing followed by freeze substitution and imaged by FIB-SEM (**Hela-1**) from the EMPIAR database (38) which we segmented for the endoplasmic reticulum (ER), nuclear envelope (NE), mitochondria, and the Golgi apparatus. A small subset of this dataset was used to train CebraNET. Secondly, we segmented a similar dataset, also containing a HeLa cell, also prepared by chemical fixation and freeze substitution and also acquired by FIB-SEM, which we used to prove the transferability of the method (**Hela-2**). Here, we segmented the ER, representing a fine-detailed organelle, as well as mitochondria, to show the quality when segmenting individual instances at a lower resolution. Thirdly, further transfer was shown with a dataset of a SARS-CoV-2-infected Calu3 cell which, in difference to the first two datasets, was chemically fixed and imaged by FIB-SEM (**Calu3-3**). We here segmented ER and mitochondria as well as the virus-induced double-membrane vesicles (DMVs). Furthermore, we used a dataset showing a macrophage that contained a parasitic vacuole. The sample was, as the previous, prepared by chemical fixation and imaged by FIB-SEM (**Macrophage-4**). Here we focused our segmentation effort on the ER of the host as well as the parasitic ER. Finally, we segmented a dataset showing *Platynereis dumerilii* larvae prepared by chemical fixation and imaged by serial block-face SEM for mitochondria in ciliated cells. We thus proved the applicability to multicellular organisms and the scalability of our approach to datasets in the size scale of tera-bytes (**Platynereis-5**).

## Results

CebraEM is a workflow that combines an efficient segmentation step for the annotation of training data (Fig. 1a and b) followed by a fully automated shallow learning-based inference approach (Fig. 1c). CebraEM’s strengths articulate around three pillars, with the aim to make it accessible to a wide community of users: (I) it avoids the necessity to intensively train a neural network, (II) the inference model is trained on annotations of the same target dataset that the user chooses to segment, thus approaching the result to the specific domain of the data, and (III) it yields a fully learned semantic instance segmentation.

**Fig. 1.**
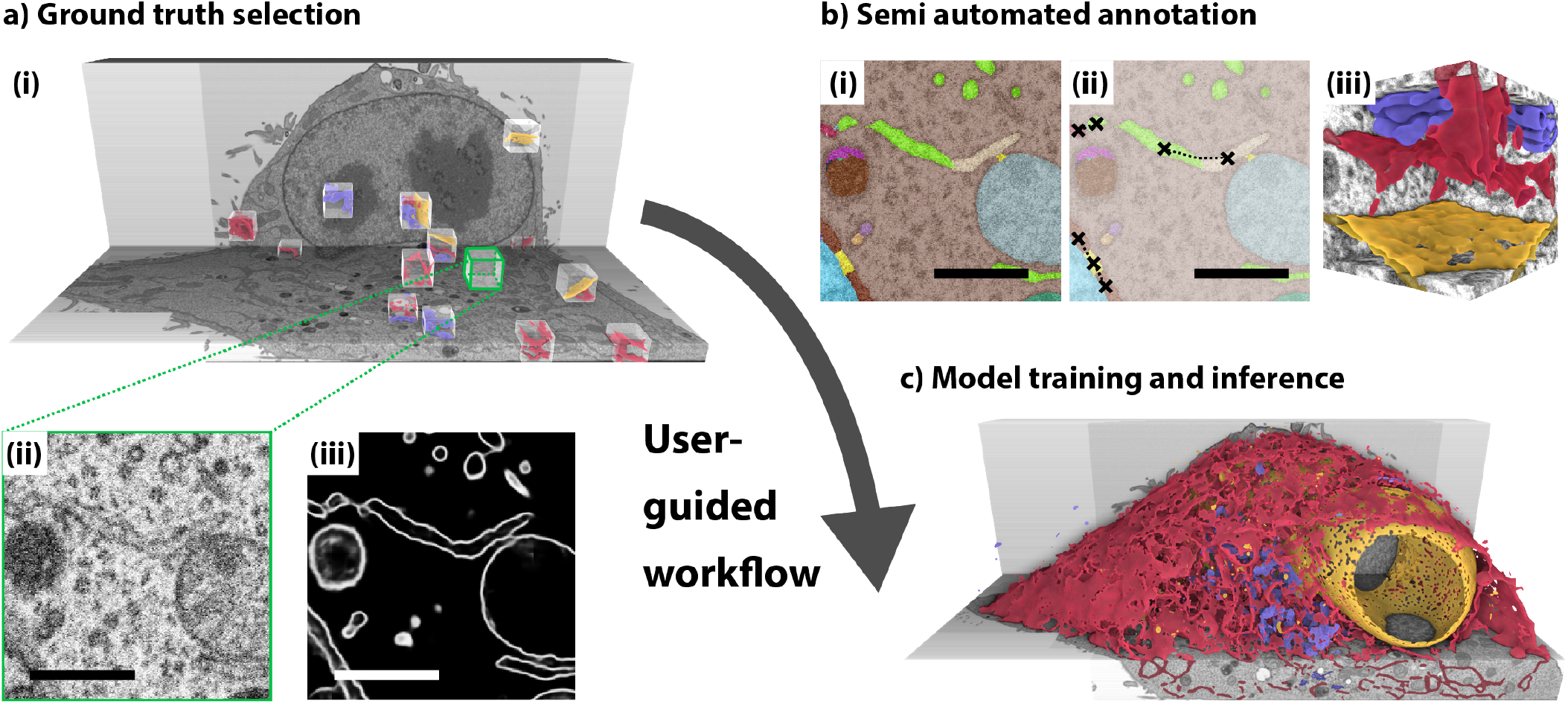
Overview of segmentation workflow. a) Selection of ground truth cubes throughout the dataset showing (i) semi-automated annotations of ER, NE, and Golgi stacks. cubes were sampled manually along the full volume and extracted for processing. (ii and iii) Each cube is passed to the membrane prediction network, enhancing the edges of membranes where organelles are present. This step will facilitate the subsequent supervoxel-based segmentation (iv). b) Semi-automated annotation of the cubes selected in the previous step, with manual correction of an initial object-wise segmentation. (i) Result of an automatic segmentation (pre-merging) which forms the starting point for manual corrections. The result is mostly over-segmented by default. (ii) The user now corrects the organelles of interest, the ER in the present case (the crossed connected by dashed lines indicate merging events to correct for an over-segmentation of the respective ER sheets). c) Within the semi-automated workflow, it is possible to annotate several organelles, like ER (yellow), nuclear envelope (yellow), and Golgi apparatus (blue) alongside by assignment of the respective organelle class after completing each individual instance. The rest of the organelles are ignored and automatically tagged as a part of the background. When all cubes are annotated, they are used to train the segmentation models for each class, which are then applied to the full dataset. Scale bars: 0.5 µm.

From a user perspective, the CebraEM workflow starts with the initialization of a CebraEM project which can be visualized using the MoBIE viewer (39). Using the viewer, the user navigates the dataset and selects suitable positions for ground truth cubes that contain the required target organelle(s) (Fig. 1a-i indicating the location of suitable cubes and Fig. 1aii showing a raw data slice). After computation of a boundary prediction and supervoxels (Fig. 1a-iii and a-iv), the user annotates the ground truth cubes using the CebraANN annotation tool to obtain a dense annotation of 6 to 12 sub-volumes of 256 × 256 × 256 voxels, each (Fig. 1a-i, Fig. 1b-i and b-ii). This task was feasible within approximately 15 to 30 minutes per organelle per sub-volume, depending on its complexity (see section “Semi-automated annotation”). This normally yielded enough ground truth for training a new model (Fig. 1c, see section “Automated segmentation step”).

Following an active learning strategy, we found that it was beneficial to iterate the training process in smaller batches. Hence, after prediction on a subset of the data, one or several new cubes were selected for annotation in locations where the prediction accuracy was low. We iterated ground truth annotation and computation of intermediate results until the prediction quality started to converge. We observed that this targeted selection of ground truth cubes can improve both the speed and quality of the entire segmentation process. As a rule of thumb, a total of 3 annotated cubes was a good starting point with an increase of 2 new cubes in each iteration. Note that this strategy should be adapted to data size, diversity within the segmentation class, and quality of the intermediate segmentation results.

CebraEM capitalizes on a readily trained boundary prediction CNN (CebraNET). We observed that, when trained for boundaries, a model generalizes well and reliably picks cellular membranes. Membranes appear very similar in most volume SEM datasets, regardless of biological or experimental variability or sample preparation variations, and thus enabled the prediction of membranes without the need for retraining on new target data (see section “Transferable membrane prediction”). The output of the CebraNET was used as a base for an initial partitioning of the volume data into a strict over-segmentation using supervoxels (Fig. 1a-ii, a-iii, and b).

The training workflow is an adapted version of the connectomics multicut pipeline (20). The multicut pipeline uses a boundary prediction specific for neuron plasma membranes, from which supervoxels are computed. Based on the super-voxels, a graph representation is created where supervoxels form the graph nodes and their local connection forms the edges. Weights are assigned to each edge by prediction of their likelihood to belong to the same object (e.g., organelle or cytoplasm) based on a set of features that score each edge (e.g., values associated with texture, shape, or gray values). The resulting weighted graph is partitioned employing a multicut algorithm that globally optimizes the edge weights within each object. By doing this, the model not only learns to connect the target organelle supervoxels but also to differentiate it from the background and adjacent objects, even if they are of the same class. Consequently, even if the segmented objects are in contact, this training step can result in a semantic instance model. After the model is trained and validated on the provided ground truth cubes, the full dataset can be segmented by inference of the model to the entire volume (Fig. 1c). At any time, from initialization to the final segmentations, the CebraEM project with its associated raw data, membrane prediction map (CebraNET), supervoxel map, and segmentation maps can be viewed using the MoBIE viewer (39).

### Transferable Membrane Prediction

As described in the introduction, one of the key aspects of segmentation is to split the dataset into separated regions that accurately delimit the organelles’ boundaries. Since the sample preparation for the volume EM dataset is generally tuned to make cell membranes visible, we trained a CNN to generate membrane predictions (CebraNET) as a basis for this initial partition.

First, a section of 512 × 512 × 64 voxels was selected from a publicly available FIB-SEM dataset (EMPIAR-10311) which contained a high number of cellular organelles. This section was densely annotated for organelles using Microscopy Image Browser (40). From this organelle segmentation, we derived object boundaries, which we defined as pixels located at a change of object label in the segmentation. Note that we masked out the inner part of organelles in the model training process.

The generated boundary ground truth, in conjunction with masked organelle interior, was used to train a concatenation of three 3D U-Nets (23) in order to predict organelle boundaries in volume SEM data (Fig. 2a). For training, we used a binary cross entropy loss applied to the output of each successive U-Net module thus comparing the output to the organelle boundary map. Additionally, we combined different types of data augmentation (including brightness and contrast variations) to achieve a better generalization of the model and successfully predict dim boundaries in the test datasets. The final model, which we called CebraNET, can be seen as a dedicated boundary predictor for volume SEM datasets, predicting plane-like structures like cellular membranes. The predicted membrane edges delimit separated regions, which enabled a good initial partitioning into supervoxels. This is a crucial step since the subsequent steps require a decisionmaking process to join those supervoxels into full objects.

**Fig. 2.**
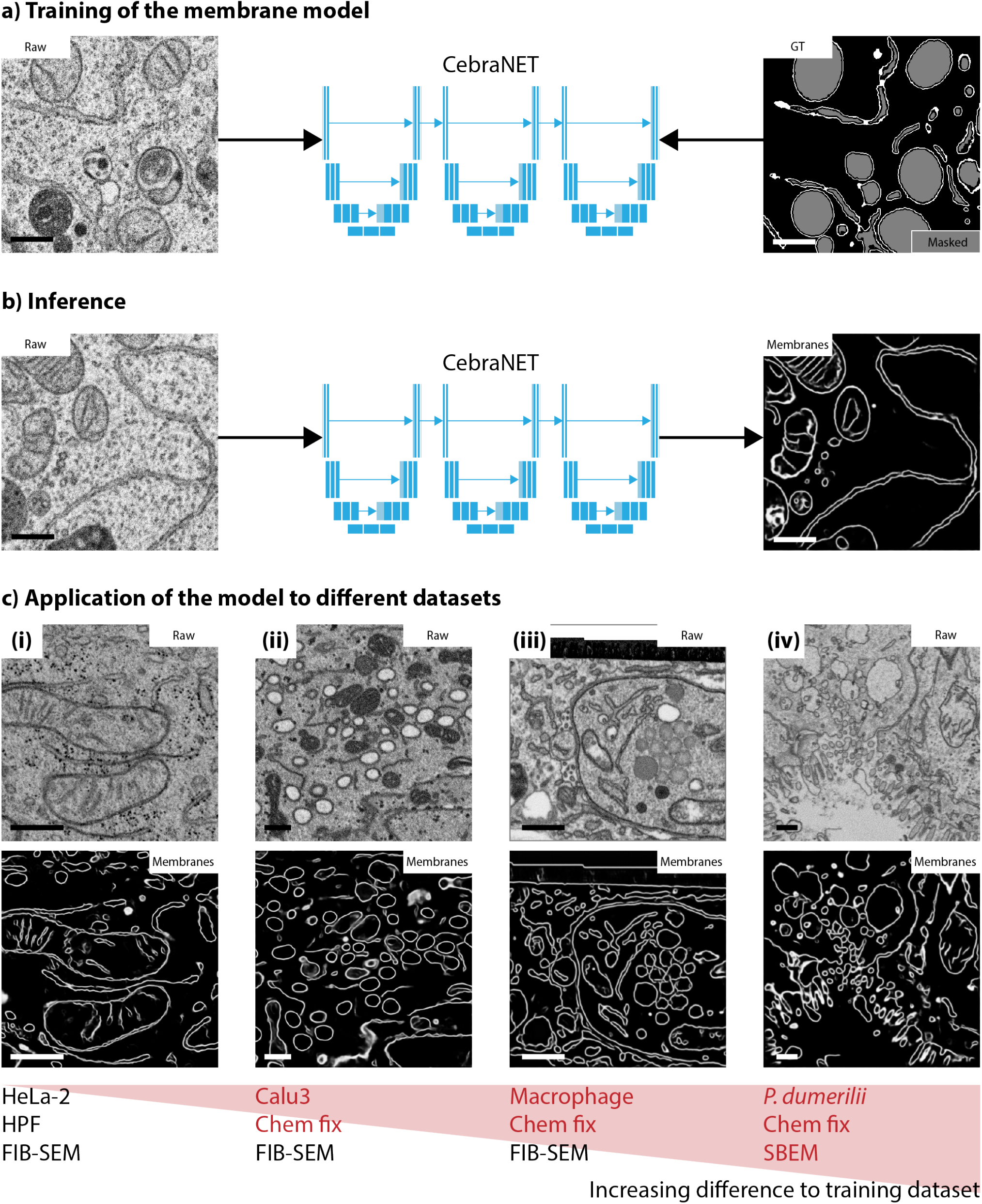
Application of our CebraNET model to a variety of different volume SEM datasets to identify membranes. a) A U-Net-based model was trained on a small subset of the **HeLa-1** dataset. The model is composed by joining three blocks (ii), each block composed of a U-net. The input of the model was the FIB-SEM data, and the output was the annotated outer membranes of organelles in form of a white mask, with the areas inside organelles masked (gray), and the background in black, ignored. The loss function was computed as the difference between masked annotations and the output of the network as a probability map of being a membrane. b) After training, the same model is applied to new raw FIB-SEM data, resulting in a prediction of almost all cellular membranes including those which have not been used during the training (e.g. within organelles such as the mitochondrial cristae). c) The inference model was successfully applied to other datasets, displayed here with an increasing difference to the original training dataset: (i) **HeLa-2**, another Hela cell, also processed using high-pressure freezing and imaged with the same microscope type, a FIB-SEM, (ii) **Calu3-3**, which is a human lung cancer cell, with different treatment, chemical fixation, and same microscope type (FIB-SEM), (iii) the **Macrophage-4** dataset, from a chemically fixed cell imaged with a FIB-SEM and, finally (iv) **Platynereis-5**, from a chemically fixed multicellular annelid, imaged with an SBF-SEM. Scale bars: 0.5 µm.

In practice, when applied to a different dataset, CebraNET was able to infer membrane-like structures, even if they were dissimilar to the original dataset (Fig. 2b and c). In the **HeLa-1** and **HeLa-2** datasets, membranes of a broad variety of organelles were successfully predicted including inner-organelle membranes, with the input data scaled to a resolution of 5 nm isotropic for the **Hela-1** dataset (Fig. 2b) and the native 4 nm resolution for the **Hela-2** dataset (Fig. 2c-i). For the **Calu-3** dataset, organelle boundaries of mitochondria and DMVs were well predicted at the native 8 nm isotropic resolution (Fig. 2c ii). However, compared to the HeLa datasets, the ER was not well reconstructed. An artificial increase of the resolution to 5 nm isotropic yielded a good boundary reconstruction such that the segmentation of ER in this dataset became feasible (Supplementary Figure S1). In contrast, the **Macrophage-4** dataset, in its native 5 nm resolution, yielded a boundary prediction with a high level of detail, such that the segmentation of the ER was feasible in its native resolution (Fig. 2c iii).

The **Platynereis-5** datasets showed the strongest anisotropy of the tested datasets as well as the lowest xy-resolution. For the segmentation of mitochondria, we mapped the voxels of 10 × 10 × 25 nm^3^ anisotropic input data to an isotropic resolution of 10 × 10 × 10 nm^3^. In this way, we improved the accuracy of the prediction for mitochondrial outer membranes and enabled the downstream steps for this dataset (Fig. 2c iv). Overall, using different input resolutions to the network appeared to be an efficient method to target boundaries of a certain thickness and distance. Close membranes, such as the outer double membranes of mitochondria and DMVs could be well predicted as a single boundary when using a low resolution (supplementary Figure S1 c-ii and d-ii). On the other side, when close membranes were required to resolve narrow organelles such as ER and Golgi apparatus, a higher resolution led to a boundary prediction of both (supplementary Figure S1 a-ii and b-ii). We argue that CebraNET learned to predict boundaries within a certain frequency range, similar to a band pass filter, where the adaptation of the frequency could be controlled by the input resolution.

### Semi-automated annotation

After the application of CebraNET to the raw data, the resulting membrane probability map is converted into supervoxels. To generate them we use an approach similar to the one described by Beier et al. (20) based on a seeded watershed algorithm. In our implementation, the input for the algorithm is a combination of the membrane prediction map (in areas where membrane probability is larger than a threshold *t*) with a distance transform computed on the thresholded membrane prediction map (where membrane probability is smaller than *t*). Then, local maxima are used as seeds, which grow into small partitions limited by the combined map. As a result, the watershed algorithm partitions the data into supervoxels (Supplementary Figure S1). In the annotation workflow, these supervoxels are shown to the annotator, whose task is to correct a segmentation on the supervoxel level to form instances of target objects, and then assign each to the respective semantic class (e.g., mitochondria class). To reduce the complexity of this manual supervoxel-level correction task, an initial instance segmentation is computed by a multicut-based graph optimization (20). The optimization step is set up similarly to the step of the automated segmentation (see section “Automated segmentation step”), with the difference that edge probabilities are defined by the average membrane probability along the edge of adjacent supervoxels. The resulting segmentation may contain over or under-segmentation and is not specific to any targeted object (Fig. 1b-i). The annotator can correct the segmentation by merging or splitting off supervoxels (Fig. 1b-ii) and review the result until all objects are considered correctly segmented. Alongside, the annotator can tag each segmented object to its respective class, hence, achieving semantic instance segmentation (Fig. 1b-iii).

We performed annotations on cubes of approximately 256 × 256 × 256 voxels. This size enabled a good trade-off between an overview of the data at hand, where the volume section is still large enough to include substantial amounts of the target structures and resulting annotation times. Annotation times lay mostly within the range of approximately 15 to 30 minutes for each dataset and varied strongly depending on the complexity of the target organelle morphology and the number of instances present in the annotated cube. Therefore, the required amount of ground truth for training a data-specific model could be generated in a few working days (Fig. 3 and supplementary tables S1.1, S2.1, S3.1, S4.1, and S5.1).

**Fig. 3.**
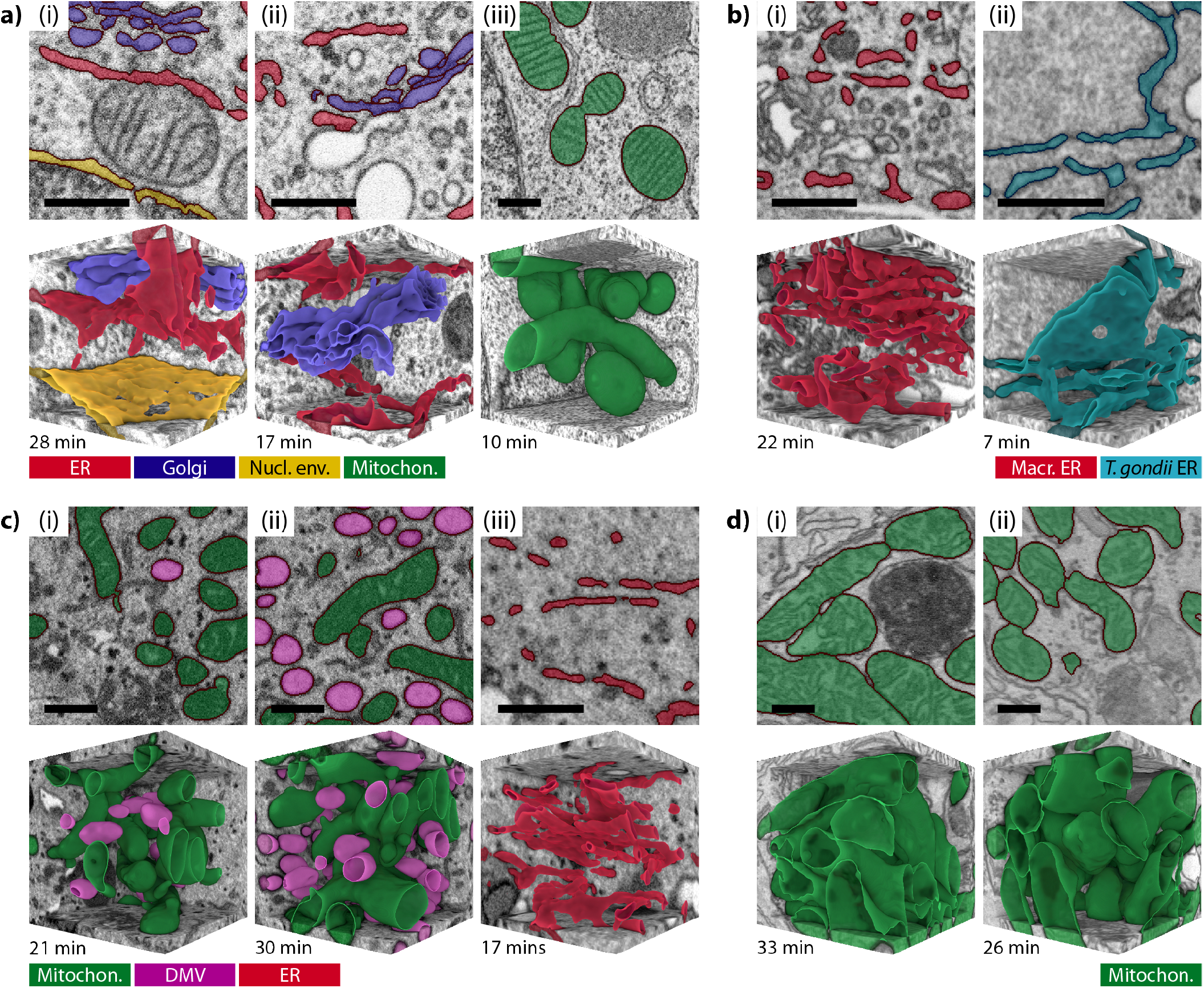
CebraANN - ground truth organelle annotation workflow: Annotation examples on 256^3^ voxel subsets, used for model training. Most annotations, even when multiple organelles were jointly annotated, could be finished in less than 30 minutes for a non-expert user of the CebraANN workflow. In every image, the time needed to complete the annotation is displayed at the bottom left of the 3D rendering. The input data volumes were scaled to isotropic voxel sizes in different resolutions to optimize annotation accuracy and annotation speed. Cubes of lower resolution (e.g., 10 nm) can be annotated much faster with respect to the sample volume they cover, however, do not yield enough information to successfully annotate more detailed organelles such as the ER or Golgi apparatus. a) **Hela-1** cell annotated (i and ii) for ER (red), Golgi apparatus (purple), and nuclear envelope (yellow), at 5 nm isotropic voxel resolution; and (iii) mitochondria (green), in this case at 10 nm isotropic voxel resolution. b) **Macrophage-4** cell annotated separately for (i) ER of the host cell (red) at 5 nm isotropic voxel resolution, and (ii) the ER of the *T. gondii* parasitic cell (dark cyan) at 4 nm isotropic voxel resolution. c) **Calu3-3** cell annotated for (i and ii) mitochondria (green) and double-membrane vesicles (DMVs, pink) at 8 nm isotropic voxel resolution, as well as for (iii) the ER (red) at 5 nm isotropic voxel resolution. d) **Platynereis-5** cells annotated for mitochondria (green) in different ciliated cells (i and ii) at a 10 nm artificial voxel size. Scale bars: 0.5 µm.

In order to maximize the efficiency of annotation, we used the most suitable resolution for each organelle. Organelles such as ER, NE, and the Golgi apparatus were annotated at a resolution of 4 to 5 nm isotropic to be able to include all their fine structural details (see supplementary figures S1a and b for an illustration using the **Calu-3** dataset). For larger organelles, such as mitochondria or double-membrane vesicles (DMVs), we used 8 or 10 nm isotropic thus increasing cellular context in computations and increasing computation speed while the supervoxels boundaries still defined the organelle instances well (supplementary figure S1c and d). In practice, within one dataset, we grouped organelles of similar size scale (e.g. ER, NE, and Golgi apparatus or mitochondria and DMVs) to be able to jointly annotate them and thus additionally increase the efficiency of the annotation process.

More specifically, the **Hela-1** dataset (native voxel size at 5 × 5 × 8 nm^3^) was annotated for ER (11 cubes), NE (6 cubes), and Golgi (6 cubes) using an up-sampled 5 nm isotropic resolution as well as mitochondria (4 cubes) using 10 nm isotropic resolution (Fig. 3a). The annotation time for the ER, NE, and Golgi in the **Hela-1** dataset was 12.8 ±6.2 (SD) minutes per cube covering a total cellular volume of 31 µm3. For mitochondria, the annotation time was 11.5 +-1.9 (SD) minutes per cube covering a total volume of 67 µm3 (Fig. 3a and Supplementary Table S1.1). For the other datasets, i.e., the **Hela-2, Calu3-3, Macrophage-4**, and **Platynereis-5** datasets, we proceeded accordingly (Fig. 3b to d and Supplementary Tables S2.1 - S5.1). Organelles of similar size scale were grouped to an equal annotation resolution (e.g., DMVs and mitochondria in the **Calu-3** dataset were annotated using 8 nm isotropic resolution). For finer structures such as the ER, we annotated at higher resolution (4 to 6 nm), thus enabling a more detailed membrane probability map that resulted in finer supervoxels (e.g., the *T. gondii* and macrophage ER were annotated using 4 and 5 nm isotropic resolutions, respectively).

To guide a user most efficiently through the process of data annotation, we implemented a plugin for the Napari viewer (supplementary Figure S7). An extracted ground truth cube containing the raw data, the pre-computed membrane prediction, and the supervoxel maps can be loaded and visualized in Napari (Fig. S7a). The pre-merging is computed as described (Fig. S7b) and then the user corrects it to form an instance segmentation containing instances of one or more organelle classes. For example, the segmentation in this layer can jointly contain instances of ER, Golgi, and mitochondria (Fig. S7c). Each completed instance is then moved to its specific semantic instance layer (one layer per organelle class) which forms the final instance segmentation maps (Fig. S7d).

### Automated segmentation step

The automated segmentation workflow that is implemented in the CebraEM package can be considered an automated version of the semiautomated workflow described above. In both workflows, an instance segmentation is generated first based on the membrane prediction and supervoxels. After this, each object is assigned to a semantic class, yielding the final semantic instance segmentation. This part is implemented using multiple components: a random forest model yields information on merging or splitting adjacent supervoxels (edge model) in conjunction with the multicut algorithm (graph-based instance segmentation algorithm based on solving a global objective) and a second random forest model, trained to determine the likelihood of a supervoxel to be part of the target organelle (semantic model).

Both random forest models, the edge and the semantic model, are trained using features derived from the annotated subsets. The feature space is set up from convolutional and statistical features extracted from the raw volume, the membrane prediction map, and the supervoxels. They comprise edge features (intensity statistics along supervoxel edges on the raw data and membrane map), region features (intensity statistics within each supervoxel from the raw data), and shape features (supervoxel shape measures). The edge model is trained using all computed features while the semantic model is trained exclusively on region and shape features (Fig. 4a-i).

**Fig. 4.**
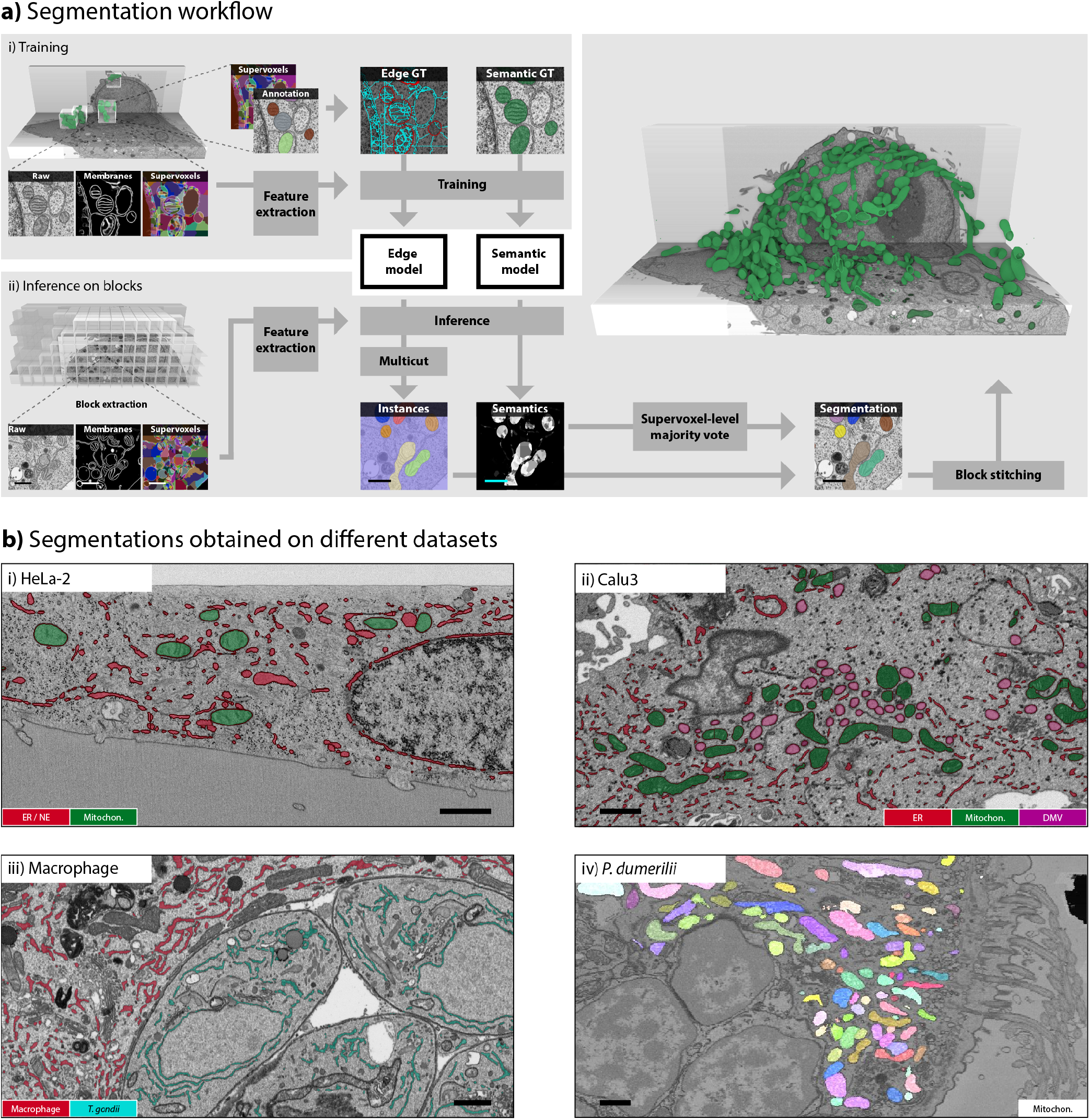
Workflow and results of the CebraEM segmentation. a) The CebraEM workflow is illustrated here for mitochondria segmentation in the **Hela-1** dataset. During training (i), randomly selected cubes are annotated on top of an initial suggested supervoxel segmentation. Once supervoxels are joined and assigned to classes, annotated cubes become input to the program, which uses them for two parallel tasks: a random forest model (edge model) uses the edge prediction of CebraNET which is thus trained to estimate the likelihood of merging two adjacent supervoxels; a second random forest model (semantic model) is trained to predict the likelihood of one supervoxel to belong to the target organelle. With both models, inference (ii) is performed block-wise for the full dataset. The edge model output is transformed into segmentation by a multicut algorithm, each component considered as an instance. The final assignment of an instance to an organelle class is decided by using the semantic model, where a supervoxel-level majority vote decides the class for each individual instance. The blocks are stitched by object overlap at the block boundaries. Scale bars: 0.5 µm b) Segmented example subsets of four different datasets: (i) **Hela-2** cell, segmented for ER (red) and mitochondria (green), (ii) **Calu3-3** cell, segmented for ER (red), mitochondria (green) and double-membrane vesicles (DMVs) (purple), (iii) **Platynereis-5** cells, segmented for mitochondria (multicolored, each color corresponds to a mitochondria instance) in ciliated cells. Scale bars: 1 µm

For inference, the full target dataset is processed as evenly spaced blocks, processed individually with a certain overlap (32 pixels in each dimension) to avoid boundary issues. First, for each block, features are extracted from raw images, membrane, and supervoxel maps as described for the model training. Second, the inference results of the edge and semantic model are obtained. The edge probabilities are converted to edge weights and passed to the multicut algorithm, which solves the multicut objective for each of the blocks, generating an instance segmentation. Finally, classes are assigned to each instance using the semantic model output. The class of each individual object is defined by a majority vote of its supervoxels, where each supervoxels votes for its most likely associated class. The results for all blocks are stitched back to the full volume by greedily merging organelle instances at block boundaries, if they intersect by more than 50 percent, thus achieving a consistent segmentation for the entire dataset (Fig. 4a-ii).

To examine the versatility of our approach we performed a segmentation of organelles for multiple datasets. In the **Hela-1** dataset, we segmented ER, Golgi apparatus, mitochondria, and nuclear envelope (Fig. 5a and b), and in the **Hela-2** dataset we segmented ER (including the nuclear envelope) and mitochondria (Fig. 4b-i). The **Calu3-3** dataset was segmented for ER, mitochondria, and SARS-Cov2-induced double-membrane vesicles (DMVs) (Fig. 4b-ii). For the **Macrophage-4** dataset, we segmented the ER from both the macrophage host and the internal parasite *T. gondii* (Fig. 4b-iii). Finally, we challenged the workflow with a very large SBF-SEM dataset, by segmenting the **Platynereis-5** dataset for densely packed mitochondria in ciliated cells (Fig. 4b-iv).

**Fig. 5.**
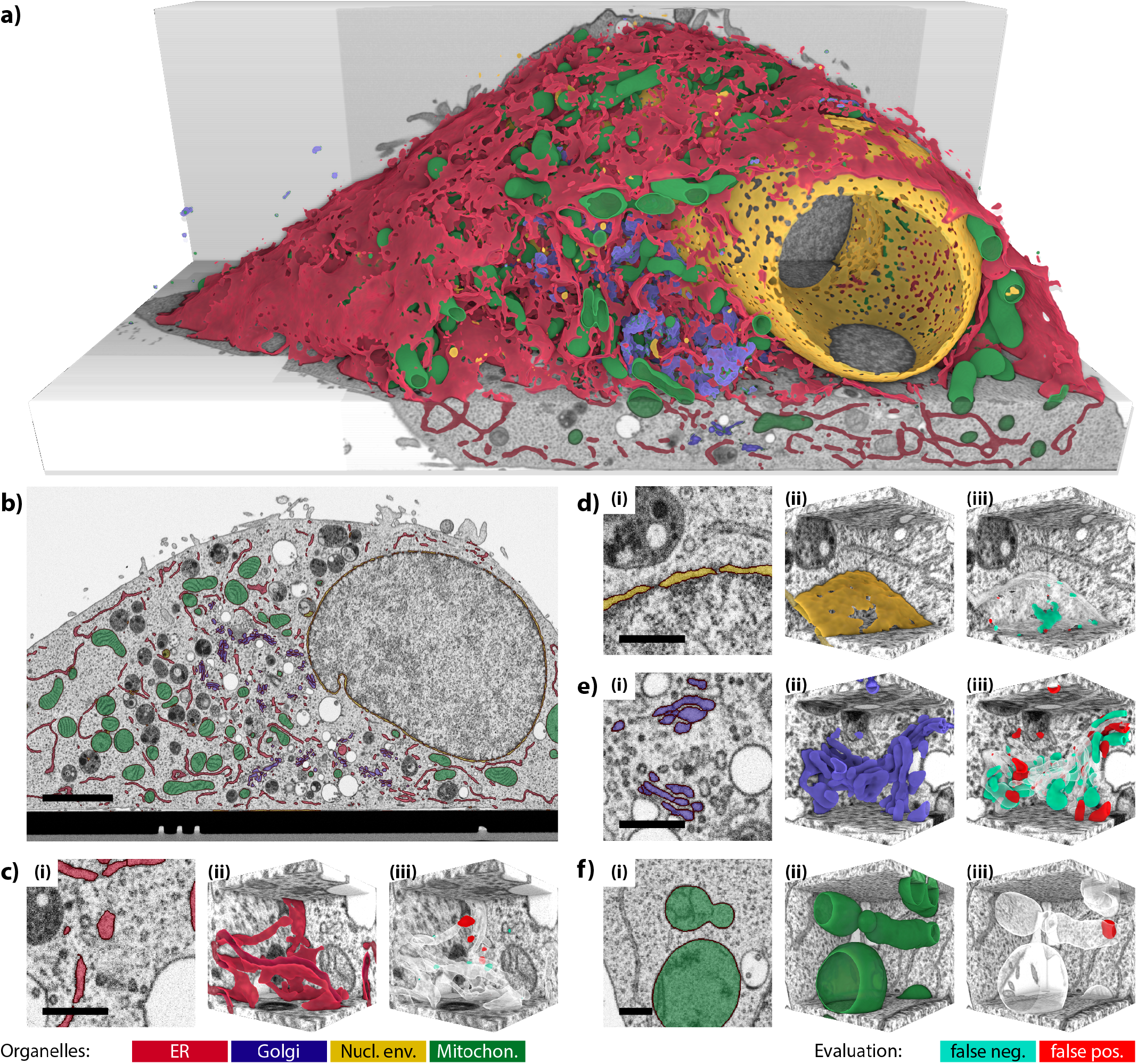
Segmentation of multiple organelles using CebraEM on the Hela-1 dataset. a) Rendering of ER (red), mitochondria (green), Golgi (blue), and nuclear envelope (yellow) segmentations of the full dataset. b) Overlay of the segmentation on an example ortho-slice of the dataset marking the organelles (same colors as in (a)). c) to f) Evaluation of the automated segmentation using validation cubes. Each panel shows, for each segmented organelle: (i) an overlay (selected 2D slice) of the automated segmentation of the FIB-SEM volume, (ii) a 3D rendering of the segmented objects, and (iii) a 3D rendering of the comparison between the manually annotated cubes and their prediction (annotated cubes used only for validation, not for training), where red areas denote false positives and cyan areas denote false negatives. c) In red, ER with an IoU of 0.99, d) in yellow, nuclear envelope with an IoU of 0.92, e) in blue, Golgi apparatus with an IoU of 0.63, and f) in green, mitochondria with an IoU of 0.93. The resulting accuracies can vary between the annotator, the location of the cubes, and the organelles segmented. As an example, in the case of e), Golgi cisternae were manually annotated whilst the automated prediction also included vesicles and parts of the ER close to it, increasing the number of false positives and false negatives in the region and contributing to a relatively low IoU and an overall impression of a high error rate. Scale bars are 2.5 µm (b) and 0.5 µm (c to f).

This automated pipeline yielded proper semantic instance segmentations that did not need further post-processing. Furthermore, for average-sized datasets typically obtained from a FIB-SEM session (e.g. 16,000 µm^3^ at 8 × 8 × 8 nm^3^ voxel size, 3 days acquisition on an off-the-shelf FIB-SEM), results were obtained within two weeks, from the generation of the ground truth on cubes at selected locations to the complete segmentation of the entire volume.

### Evaluation of accuracy

To evaluate the quality of the automated segmentations, we annotated additional validation cubes for each dataset and organelle using the CebraANN workflow. For each dataset, we selected extra cubes at random locations. We ensured that each cube that was used in the validation contained the desired target organelle and did not overlap with any of the training cubes. Then, we compared the automated segmentation result with the validation data using the ratio of correctly segmented areas against the union of true positive and false positive segmented areas (also known as IoU, Intersection over Union) (Table 1 For example, for the **Hela-1** dataset, our segmentations were validated with 8 cubes containing ER, 3 cubes containing NE, 3 cubes containing Golgi apparatus, and 3 cubes that contained mitochondria. Since ER, NE, as well as Golgi, were annotated at the same resolution level (5 nm isotropic), we used the same cube for multiple organelles when applicable. The segmentations for mitochondria provided an IoU of 0.96 ±0.04 (SD) (Fig. 5f-iii) and an ER of 0.93 ±0.08 (SD) (Fig. 5c iii). The other organelles had values of 0.73 ±0.17 (SD) (Fig. 5d-iii) for NE and 0.65 ±0.04 (SD) (Figure 5e-iii) for the Golgi apparatus. Note that in the Golgi apparatus, most IoU errors originated from false negatives around a well-detected inner core of the respective Golgi stack, indicating ambiguities in the assignment of objects as either vesicles or part of the Golgi stack. However, all Golgi stacks present in the full volume were found by the prediction (Table 1, row **Hela-1**).

**Table 1.** Annotation quality of the automated segmentation workflow by means of IoU for the **Hela-1, Hela-2, Calu-3, Macrophage-4** (containing ER from the *T. gondii* parasites as well as the macrophage cell), and **Platynereis-5** datasets.

|  | Mito | ER | NE | Golgi | DMV |
| --- | --- | --- | --- | --- | --- |
| **Hela-1** | 0.96 ± 0.04 | 0.93 ± 0.08 | 0.73 ± 0.17 | 0.65 ± 0.04 | - |
| **Hela-2** | 0.89 ± 0.12 | 0.73 ± 0.09 | - | - | - |
| **Calu3-3** | 0.78 ± 0.20 | 0.78 ± 0.15 | - | - | 0.99 ± 0.01 |
| **Macrophage-4: macrophage** | - | 0.93 ± 0.06 | - | - | - |
| **Macrophage-4: *T. gondii*** | - | 0.84 ± 0.04 | - | - | - |
| **Platynereis-5** | 0.82 ± 0.14 | - | - | - | - |

The segmentations of the other four datasets (**Hela-2, Calu3-3, Macrophage-4**, and Platinereys-5) were evaluated in the same way. We used a minimum of three validation cubes per organelle to obtain reliable evaluation metrics. Our approach achieved IoU scores of 0.7 or higher, mostly depending on the difficulty of the respective organelle. For example, ER in the **Calu3-3** dataset yielded a final IoU of 0.78 ±0.2 (SD) while DMVs in the same dataset achieved an IoU of 0.99 ±0.01. In many cases, we were able to achieve segmentation of IoU > 0.8 after two iterations of annotations (supplementary tables S6 to S10). The first iteration establishes an initial set of ground truth cubes and the second iteration adds more information at locations where the initial model underperformed. This was the case for the segmentations of both ER types (macrophage and *T. gondii*) in the macrophage dataset with an IoU of 0.84 ±0.04 (SD) and 0.93 ±0.06 (SD), respectively, as well as for the segmentation of mitochondria in the **Platynereis-5** dataset with an IoU of 0.82 +-0.14 (Table 1).

Since the data partitioning was based on a 3D boundary prediction, the object boundaries of the generated organelle instances showed smooth and plausible appearances in all three spatial dimensions. However, the segmentation quality still depended on image quality as well as some morphological aspects of the target organelles. Small objects (only a few pixels in width) were often not resolvable, as there was no significant space between the predicted membranes to form suitable supervoxels in the downstream processing. This situation arose, for example, for narrow stretches in the ER as found in the **Hela-2** dataset (Supplementary Fig. S8). In other cases, two significant membranes could be recovered by increasing the resolution artificially (e.g. from 8 nm to 5 nm as used in the **Calu-3** dataset, supplementary Figures S1b and c). The same effect, i.e. a missing space in-between two membranes, could be observed at inter-organelle cytoplasmic space, observed for example, at some mitochondria-ER contact sites (Fig. 5b, **Hela-1**) or adjacent Golgi cisternae (Fig. 4b-i). Consequently, thin objects or thin areas of larger structures were merged into the surrounding region, usually the cytoplasm, reducing the overall accuracy slightly.

## Discussion

Segmentation of volume SEM datasets is a challenging task with data-dependent difficulties originating from sample type, sample preparation, biological perturbations (e.g., knock-downs, drug effects, or virus infections), microscope hardware, image acquisition parameters, and system stability. With such a diversity of variables affecting the final data appearance, training a generic organelle segmentation model or using unsupervised approaches still remains a difficult task.

With CebraEM, we followed a strategy that prioritizes the supervised training of shallow learning models on the target data, whilst paying specific attention to the time budget necessary to achieve a satisfying semantic instance segmentation. As a result, complex segmentation tasks are achieved single-handedly within a reasonable amount of time (a few working days), even on dense volumes.

The basis for CebraEM’s efficiency and generalization to the broad variability of volume SEM datasets is our readily trained membrane model, CebraNET. The CebraNET membrane model can be adapted to the detail level of the respective target organelle by variation of the pixel size of its input, which thus forms the input pixel size to the CebraEM workflow. Within this work, we are showing CebraEMs segmentation capabilities within a resolution range from 4 to 10 nm, yielding a high level of complexity in organelles such as the ER, NE, or the Golgi apparatus, while increasing efficiency for organelles where a high-resolution level is not required (e.g. mitochondria or DMVs). We additionally demonstrate the successful application of CebraEM to multiple datasets originating from different imaging modalities (FIB-SEM and SBF-SEM), sample processing techniques (high-pressure freezing in conjunction with freeze substitution or chemical fixation), and sample types.

Most recently available segmentation approaches follow the strategy of initially creating a deep semantic model which outputs a class probability for each pixel. To then obtain individual objects of each class (i.e. a semantic instance segmentation), adapted post-processing is usually required for each individual organelle, which not only requires computation time but also computation expertise for its development and parameter tweaking. Contrary to this, our strategy, implemented in CebraEM, was to create a supervised model that concurrently yields both the semantic and the instance information and thus generates a fully learned semantic instance segmentation without the requirement for adapted and costly post-processing.

In contrast to deep semantic models, the CebraEM approach does not predict organelle classes on a pixel level as objects are confined to the supervoxels generated within the workflow. In some cases, thin objects, such as very narrow ER stretches or small cytoplasmic spaces between adjacent organelles, collapse the membrane model and thus inhibit supervoxel generation. These areas then are not detectable even in the manually guided CebraANN workflow. However, we believe, for a general-purpose tool and with the goal of maximum efficiency, this is an acceptable cost. Also, a subsequent deep learning-based approach to detect and fix such instances could be implemented.

The entire CebraEM workflow is available as an open-source package on GitHub (github.com/jhennies/CebraEM), implemented in a script-based fashion for usage on a workstation or cluster setup. The embedded annotation tool CebraANN is a graphical user interface implemented as a plugin for the image analysis software Napari. Overall, the proposed CebraEM workflow is designed to enable segmentation in large datasets without the requirement for extensive expertise in machine learning while reducing the amount of manual input. We have the conviction that CebraEM will contribute to generating faster and better 3D models for the volume SEM community. Additionally, the annotation tool CebraANN in conjunction with CebraNET could be used for other applications including the efficient generation of ground truth for deep learning models or the segmentation of entire small datasets.

## Methods

### Code availability

The code is available on GitHub (github.com/jhennies/CebraEM) with an MIT open-source license.

### Datasets

Datasets used within this publication: **Hela-1, Hela-2, Calu3-3, Macrophage-4**, and **Platynereis-5** which cover different sample preparation methods, data acquisition methods, and raw data resolutions (Table 2).

**Table 2.**
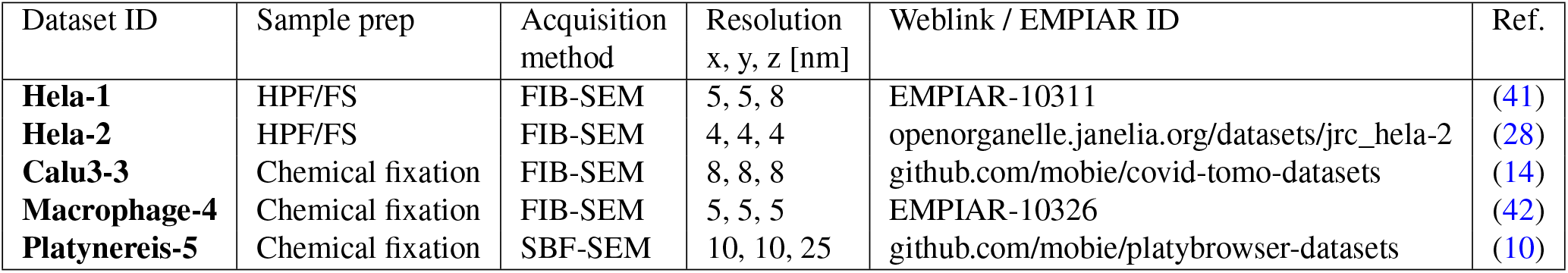

| Dataset ID | Sample prep | Acquisition method | Resolution x, y, z [nm] | Weblink / EMPIAR ID | Ref. |
| --- | --- | --- | --- | --- | --- |
| **Hela-1** | HPF/FS | FIB-SEM | 5, 5, 8 | EMPIAR-10311 | (41) |
| **Hela-2** | HPF/FS | FIB-SEM | 4, 4, 4 | openorganelle.janelia.org/datasets/jrc_hela-2 | (28) |
| **Calu3-3** | Chemical fixation | FIB-SEM | 8, 8, 8 | github.com/mobie/covid-tomo-datasets | (14) |
| **Macrophage-4** | Chemical fixation | FIB-SEM | 5, 5, 5 | EMPIAR-10326 | (42) |
| **Platynereis-5** | Chemical fixation | SBF-SEM | 10, 10, 25 | github.com/mobie/platybrowser-datasets | (10) |

### Data visualization and data storage

The big-data-viewer-based MoBIE viewer (39) was used to inspect large datasets. MoBIE enables image pyramid-based lazy loading of the data and scales to arbitrary data size. Additionally, it can display multiple datasets such as raw data, processing steps, and final segmentations in an overlay representation. We implemented CebraEM projects in a MoBIE-readable format such that they can be directly viewed without any conversion requirements. Before initialization of the CebraEM project, the raw data has to be converted to the n5-based BigDataViewer (43) format for which CebraEM ships a conversion script. All computational results (membrane prediction, supervoxels, and segmentation maps) are, on the fly, saved into the project structure in the required n5-based BigDataViewer format which enables parallel writing for increased efficiency of data I/O. Already computed subsets can then be visualized in the MoBIE viewer already before the completion of the computation.

### Preprocessing

We use AMST (41) to align each of the volume image stacks. If the data showed a varying gray value distribution over one or more data axes, we corrected the histogram along the respective dimension to ensure a homogeneous gray value distribution throughout the entire dataset. As an example, gradients in the **HeLa-2** dataset were corrected along the x- and z-axes. Before feeding the data into the CebraEM workflow, we normalized the data by histogram stretching converting the 2-percentile to 0 and the 98-percentile to 255 and subsequent conversion to an 8-bit integer data type. The quantiles for normalization were generally confined to regions containing the sample, thus excluding the effect of differing amounts of empty resin within each slice. The masks were obtained by a rough thresholding-based segmentation performed in Microscopy Image Browser (40). As in most cases for cellular volume SEM datasets, a significant amount of the data contains empty resin or non-imaged background, the masks used for data normalization were also used to limit computation within the subsequent CebraEM workflow to the areas containing the target cell(s). In the case of the Platynereis-5 dataset, the mask covering ciliated cells was obtained from a previously published cell segmentation and classification (10).

### CebraNET architecture and training

The CebraNET is set up based on the 3D U-Net architecture (23). We set up a concatenation of three 3D U-Net modules consecutively where the last convolution layer of each module forms the input to the subsequent U-Net module. Each of the three U-Net modules has its sigmoid output layer with binary cross entropy loss. All the U-Net modules are trained as one big network such that the three output layers are treated as three independent output channels.

The training was performed with a single 256 × 256 × 64 pixels subset of the **HeLa-1** dataset which was manually selected and contained multiple types of intracellular membranes (originating, for example, from mitochondria, the ER, or endosomes) (Fig. 2a). This ground truth data were annotated for the outer organelle membranes using Microscopy Image Browser, by annotating the present organelles and defining the object boundaries as outer membranes (this saved time as opposed to the annotation of the membranes themselves). To avoid passing contradictory information to the network (e.g., in the form of mitochondria cristae), we masked the insides of the organelles (Fig. 2b) A set for validation of 256 × 256 × 256 pixels from the **HeLa-1** dataset was annotated in the same manner.

For the model training, cubes of 64 × 64 × 64 pixels were extracted at evenly spaced positions in the ground truth cubes (spacing of 32 pixels in all dimensions) thus forming the training data for each epoch. The cubes were shuffled, augmented, and fed into the network with a batch size of 2. Augmentation was performed using affine transformations, additive noise (Gaussian), anisotropic smoothing (anisotropic Gaussian filter with random orientation), random displacement of z-slices to mimic slight misalignments as well as brightness and contrast alterations. Note that the data scaling as part of the affine transformations was performed in a range of 0.8 to 1.2 in order to make the model robust with respect to differences in membrane thicknesses as well as to differing input resolutions.

The network was trained for 200 epochs and model states with good validation scores were saved. After the training procedure candidate models with the highest validation score were examined qualitatively on their capability to predict membranes on different datasets and the best-performing model was selected. The model we publish alongside this paper was trained for 97 epochs. The model is additionally available within the Bioimage Model Zoo (44), at bioimage.io with doi ID: 10.5281/zenodo.7274275.

The inference was performed with a batch size of 64 × 64 × 64 pixels and an overlap of 32 pixels. To avoid too much overhead, especially on compute clusters, we process the full dataset in blocks (256^3^ or 384^3^ voxels), from which the batches are then extracted and written to a respective result volume after computation.

### Supervoxels

For the computation of supervoxels, we use an approach that we adapted from Beier et al. (20) with the basic principle of binarizing the membrane prediction map with a threshold *t*, computing a signed distance transform on the binarized map (positive values where the probabilities are > t and negative values where the probabilities are < *t*) and using this signed distance transform as the input to a seeded watershed. We adapted this approach by using the distance transform where the probabilities were smaller than t while maintaining the probability values where they were greater than *t*. Within this setting, the threshold t could be chosen close to zero (usually *t* = 0.05 showed good results), such that most of the value range of the membrane probability map was used to generate supervoxels (Supplementary Fig. S1).

### Multicut

We performed multicut computations using the elf package (https://github.com/constantinpape/elf) which implements the Multicut workflow introduced by Beier et al. (20). For the annotation workflow, we used a non-learned version of the Multicut workflow (45), where edge weights for optimizing the graph partitioning are directly derived from the intensity values of the membrane prediction map at the respective supervoxel edges (yielding a cumulative boundary probability over each supervoxel edge). For automated segmentations, we used learned edge probabilities which were computed by Random Forest classification (see section Automated segmentation). In both cases, the probabilities are converted to edge weights by 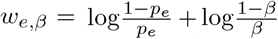 such that probabilities *p*_*e*_ *<* 1*− β* yield positive values, representing attractive edges (adjacent supervoxels should be merged), and probabilities *p*_*e*_ > 1 − *β* yield negative values representing repulsive edges (adjacent supervoxels should be split).

By manipulating the value of the beta parameter and comparing the resulting model performance, one can determine the optimal value for the parameter. We set an initial value of the parameter at 0.7 and test the result in the validation set. Then, through a process of incrementally increasing or decreasing this value by 0.05, a determination can be made as to whether a higher or lower value is more advantageous for the given segmentation result. The decision can be done by following the accuracy of IoU in a new annotated cube (a validation cube) or by visual inspection of the result. This iterative process allows for the fine-tuning of the segmentation result, ultimately leading to improved quality when applied to the rest of the dataset.

### Annotation Workflow

The annotation workflow implemented in the CebraANN package is performed on and thus optimized for, 256 voxel cubes extracted from the full dataset. The extraction of the cubes is implemented within the workflow for the specific locations (if not performed beforehand).

CebraANN is a Napari plugin that uses raw data, membrane prediction, and supervoxels. The raw data is used exclusively for guidance and display purposes. The user follows the three main annotation steps: (i) pre-merging, (ii) instance segmentation, and (iii) semantic assignment. In (i) The pre-merging is performed entirely automatically using a Multicut algorithm to globally optimize the supervoxel graph representation based on graph edge weights computed by averaging the membrane prediction at the respective supervoxel edges. The resulting objects thus confine to the predicted boundaries in the membrane prediction; however, they are neither specific for certain organelles, nor necessarily contain full objects. Depending on the preference of the user, a parameter can be set to control the amount of merging within this step.

In (ii), instance segmentation, we aim to segment all instances of organelles of interest (one or more classes). The result is generated exclusively on the supervoxel level. The user manually alters the pre-merging segmentation by merging objects, adding individual supervoxels to an object (thus subtracting individual supervoxels from the adjacent object), or creating new objects. And in (iii), semantic assignment is performed by moving a complete object to the respective semantic layer (different semantic layers for the required classes can be added on demand) by selecting its location when the respective semantic layer is active. Objects moved to a semantic layer are locked in further modification by the instance segmentation but can be moved back to the instance segmentation step if required. Consequently, steps (ii) and (iii) can be performed until all target instances are readily segmented.

### Automated segmentation

We performed the automated segmentation based on a supervised learning approach using the ground truth cubes generated by the annotation workflow to train two random forest classifiers for instance and semantic segmentations, respectively. The instance classifier was trained using statistical features along the edges accumulated over filtered representations of the raw data and membrane probability maps (Gaussian smoothing, Laplacian of Gaussian, and Hessian eigenvalues, each with Gaussian sigmas of 1.6, 4.2 and 8.3) as well as differences of statistical features within adjacent supervoxels over the unfiltered raw data (statistics include the number of voxels, kurtosis, minimum and maximum gray values, quantiles, region radii, skewness, sum, and variance), thus spanning a feature vector for each supervoxel edge. The true output for each edge was determined from a dense segmentation such that an edge should be merged if the corresponding two supervoxels belong to the same object (attractive edge) or split if the corresponding two supervoxels belong to different objects (repulsive edge). The semantic classifier was trained using statistical features obtained from the supervoxels and predicts which class (target organelle or background) the respective supervoxel most likely belongs to. Both semantic and instance segmentation is combined by first generating an instance segmentation from the output of the instance classifier (see section Multicut) and then performing a supervoxel-level majority vote within the resulting instances to assign a class to each instance.

The random forest classifiers are used from the scikitlearn (46) python package and set up of 200 trees with a maximum depth of 50 for both the semantic and the instance classifiers. These parameters were determined empirically to yield both good results while keeping the computation cost as low as possible.

### Block-wise computation and stitching

For the automated processing which is generally performed on the full datasets, the data is processed in blocks, individually for the three distinct sub-steps of the pipeline: (i) Computation of membrane prediction, (ii) computation of supervoxels and (iii) computation of a segmentation. The block size can be varied depending on the available resources, however, we generally use a block spacing of 3843 voxels. Due to processing with an overlap to mitigate boundary effects, the processed blocks are actually larger: We use overlaps of 32 pixels for the membrane prediction and 128 pixels for the supervoxels and segmentations.

The resulting computations are stitched back to respective full volumes. For each block of the membrane prediction, the overlap region is removed and the remaining inner block is written to the resulting volume. For the supervoxels, the current maximum supervoxel ID is monitored and added to each newly written block such that unique supervoxel IDs throughout the final dataset are ensured. For the segmentation, we perform more complex stitching to ensure subsections of objects that span one or more block boundaries are assigned the same object ID. Object correspondence identification is performed by IoU of objects at block boundary planes (after cropping the overlap). If objects at these block boundaries overlap more than 50 % they are considered the same object and thus assigned a common ID.

### Evaluation of accuracy

For evaluation of the accuracies of the automated segmentation, we compared subsets of the segmentation to respective manual annotations using the Intersection over Union (IoU), or Jaccard index, computed by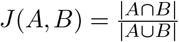, where A denotes the automated segmentation and B the manual annotation. Furthermore, *A B* is the intersecting area of segmentation and annotation, while *A B* denotes the total area spanned by both. To avoid a bias towards manually selected regions, we used a random generator to suggest positions. At a position, a cube of 256 × 256 × 256 voxels was used for validation if it (i) overlapped with the target area, i.e. within the cell or tissue, with a minimum of 75 percent, (ii) contained at least one of the target structures (i.e. target organelles) in a substantial amount and did not overlap with any of the respective ground truth cubes. Each cube was annotated for at least one of the target organelles using the CebraANN workflow.

### Data visualization and 3D rendering

To visualize and browse full EM datasets including membrane prediction, supervoxel maps, and segmentations, we used the MoBIE viewer (39). A CebraEM project is an extension of a MoBIE project such that it can be loaded directly into the viewer where each desired volume can be selected for visualization and segmentations or supervoxels can be viewed as an overlay over the respective EM data volume or membrane prediction map.

For the 3D rendering of segmentation results, we used Drishti (47). In order to enable the rendering of full datasets, we downsampled segmentation maps by a factor of 6 (with respect to the annotation resolution). Additionally, the segmentation maps were binarized (value of 255 for all instances of the target organelles and 0 for background) and smoothed with a Gaussian kernel of sigma = 0.8 in order to generate smooth object boundaries in the final rendering. Raw data was scaled to the same isotropic resolution as the segmentation maps, however not smoothed. Within Drishti, the EM data as well as segmentations were loaded as multiple volumes and the transfer functions were set such that each organelle was rendered in a different color and the raw data maintained its native grayscale colormap. To generate the cut-outs in the renderings, we added “bricks” wherein the transfer function of the EM data was deactivated. The renders of the ground truth cubes were generated in the same way with the difference that segmentations were used in their annotation resolution without further downsampling, while the raw data was scaled to match. Note that due to the binarization of the segmentations, the instance segmentation component is not color-coded in the 3D renderings. An exception is the **Platynereis-5** dataset, where we assigned random RGB colors to each mitochondria instance before loading the respective volumes into Drishti.

## Abbreviations

EM: Electron Microscopy
SEM: Scanning EM
FIB-SEM: Focused Ion-Beam SEM
SBF-SEM: Serial Block-face SEM
CNN: Convolutional Neural Network
IoU: Intersection over Union
ER: Endoplasmic reticulum
NE: Nuclear envelope

## ACKNOWLEDGEMENTS

This work was funded by the Deutsche Forschungsgemeinschaft (DFG, German Research Foundation) – Projectnumber 240245660 – SFB 1129 (project Z2)

## Supplementary Note 1: Effect of the annotation resolution

**Supplementary Fig. S6.**
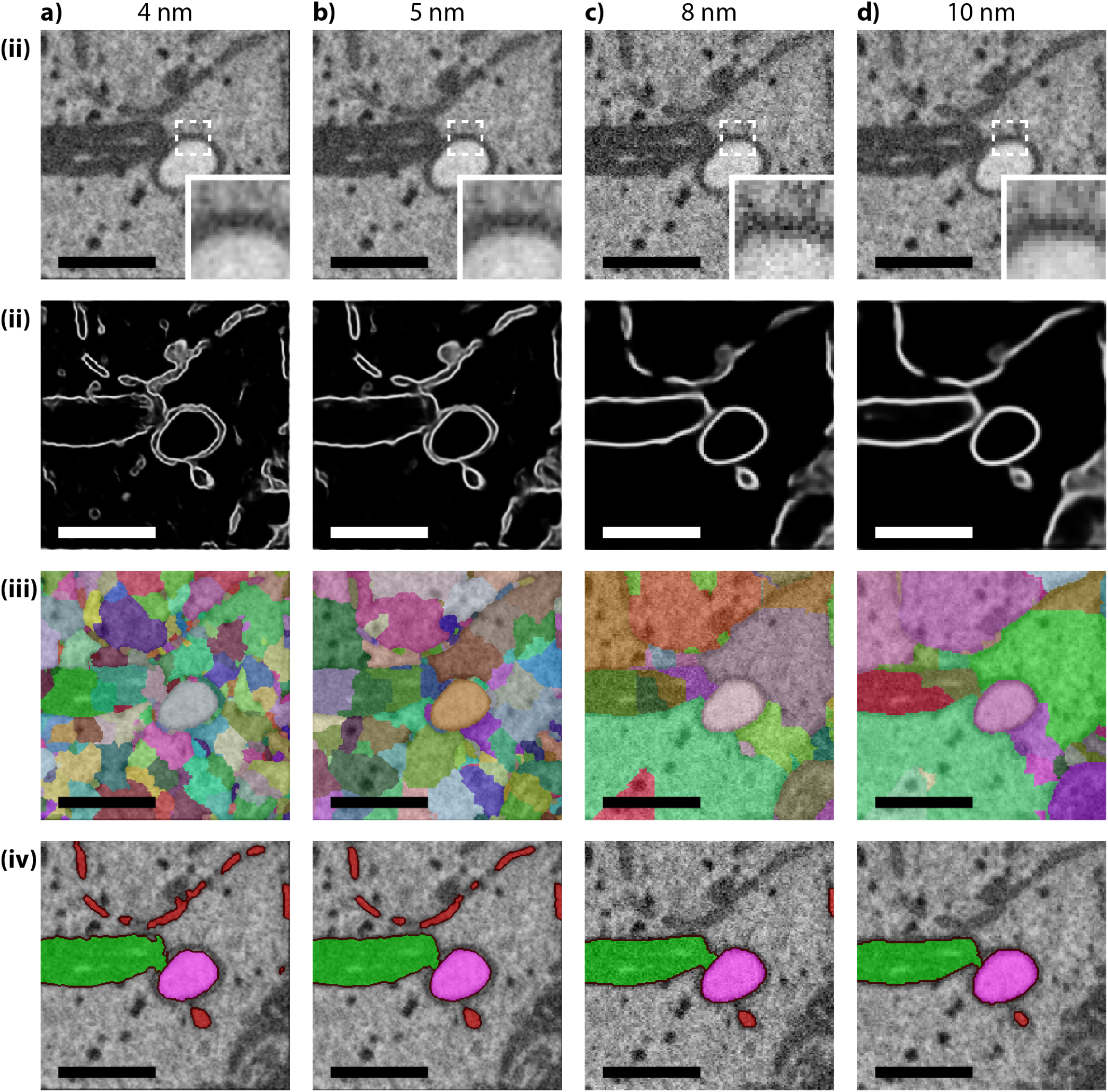
Effect of different annotation resolutions on the downstream processing illustrated with the Calu-3 dataset. The columns show the different isotropic annotation resolutions with a) 4 nm, b) 5 nm, c) 8 nm, and d) 10 nm. Note that the dataset resolution was 8 nm isotropic and is thus represented in column (c). In each column, the rows show (i) scaled input (the inset is a close-up of the area highlighted by the dashed box), (ii) boundary prediction, (iii) supervoxels, and (iv) a segmentation for ER (red), mitochondria (green) and DMVs (magenta) obtained by CebraANN. Note that the ER sheets which could be segmented using the higher annotation resolutions, 4 and 5 nm (a-iv) and (b-iv), were not resolved well in the boundary prediction of the lower resolutions, 8 and 10 nm (c-ii) and (d-ii). Here, supervoxels were only computed on rare occasions (c-iii and d-iv), such that most of the ER stretches could not be segmented using CebraANN (c-iv and d-iv). For the ER segmentation of the entire dataset, we consequently chose an annotation resolution of 5 nm. The higher 4 nm resolution did not yield further benefits and the segmented object started to appear fuzzy at the edges (a-iv). Scale bars: 0.5 µm.

## Supplementary Note 2: Hela-1

**Supplementary Table S1.3.**
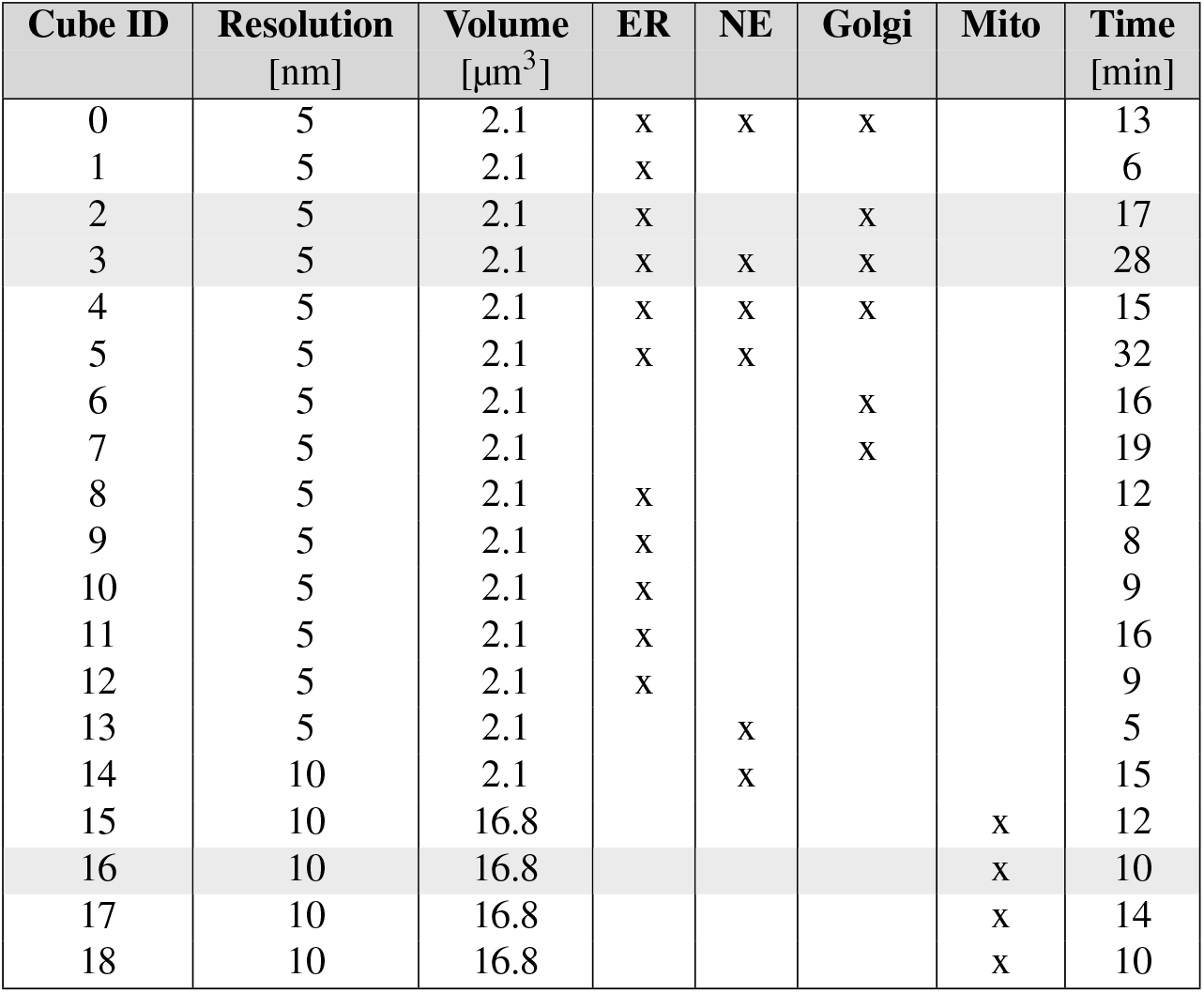
Annotated volumes and annotation time for each cube in Hela-1. Subset annotation of the **Hela-1** dataset for ER, NE, Golgi, and mitochondria (Mito). The columns refer to the cube ID, annotation resolution, cellular volume, the respective annotated organelles, and annotation time for each of the annotated subsets. The average time for annotating a cube was in the order of 14 min, depending mostly on the volume of organelle present on the cube and how many different classes were annotated on the same cube. The total volumes or annotated cubes were 23.1 µm^3^ for the ER, 12.6 µm^3^ for the NE and Golgi, each, and 67.2 µm^3^ for the mitochondria while the total cellular area was approximately 1800 µm^3^.

| Cube ID | Resolution [nm] | Volume [$\mu\text{m}^3$] | ER | NE | Golgi | Mito | Time [min] |
| --- | --- | --- | --- | --- | --- | --- | --- |
| 0 | 5 | 2.1 | x | x | x |  | 13 |
| 1 | 5 | 2.1 | x |  |  |  | 6 |
| 2 | 5 | 2.1 | x |  | x |  | 17 |
| 3 | 5 | 2.1 | x | x | x |  | 28 |
| 4 | 5 | 2.1 | x | x | x |  | 15 |
| 5 | 5 | 2.1 | x | x |  |  | 32 |
| 6 | 5 | 2.1 |  |  | x |  | 16 |
| 7 | 5 | 2.1 |  |  | x |  | 19 |
| 8 | 5 | 2.1 | x |  |  |  | 12 |
| 9 | 5 | 2.1 | x |  |  |  | 8 |
| 10 | 5 | 2.1 | x |  |  |  | 9 |
| 11 | 5 | 2.1 | x |  |  |  | 16 |
| 12 | 5 | 2.1 | x |  |  |  | 9 |
| 13 | 5 | 2.1 |  | x |  |  | 5 |
| 14 | 10 | 2.1 |  | x |  |  | 15 |
| 15 | 10 | 16.8 |  |  |  | x | 12 |
| 16 | 10 | 16.8 |  |  |  | x | 10 |
| 17 | 10 | 16.8 |  |  |  | x | 14 |
| 18 | 10 | 16.8 |  |  |  | x | 10 |

**Supplementary Table S1.4.** Evaluation of segmentation scores (IoU) for HeLa-1. For analysis, segmentations were computed with different values for the parameter β (data columns). The analysis was performed for four organelles: a) ER, b) nuclear envelope, c) Golgi, and d) mitochondria. For the ER, we performed three iterations (column It-ID) with six initial ground truth cubes for training, eight in the second, and 11 in the final iteration (column n_GT). The IoU score was computed in 8 independent annotated cubes, not used for training, for the ER, and 3 cubes for NE, Golgi, and mitochondria, each. The IoU score increased from 0.64 to 0.93 from the initial to the second iteration (It-ID 0 to it-ID 1). We observed no increase in IoU when adding more ground truth after the third iteration (It-ID 2). The highlighted cells indicate the value of the beta used for the final inference on the full datasets. Note that the final decision of which beta value was chosen was not based only on the IoU result but also on the visual inspection of the output. For NE, Golgi, and mitochondria we performed two iterations each, while mostly being able to increase the segmentation quality by adding two to three cubes of ground truth.

|  | beta: |  |  | 0.5 |  | 0.55 |  | 0.6 |  | 0.65 |  | 0.7 |  | 0.75 |  | 0.8 |  |
| --- | --- | --- | --- | --- | --- | --- | --- | --- | --- | --- | --- | --- | --- | --- | --- | --- | --- |
|  |  | It-ID | n_GT | mean | s.d. | mean | s.d. | mean | s.d. | mean | s.d. | mean | s.d. | mean | s.d. | mean | s.d. |
| a) | ER | 0 | 6 | **0.04** | 0.06 | **0.14** | 0.25 | **0.38** | 0.33 | **0.55** | 0.37 | **0.64** | 0.40 | **0.68** | 0.41 | **0.88** | 0.08 |
|  |  | 1 | 8 | **0.52** | 0.42 | **0.72** | 0.34 | **0.89** | 0.15 | **0.92** | 0.09 | **0.93** | 0.07 | **0.91** | 0.07 | **0.81** | 0.14 |
|  |  | 2 | 11 | **0.72** | 0.36 | **0.88** | 0.17 | **0.90** | 0.16 | **0.93** | 0.08 | **0.94** | 0.06 | **0.92** | 0.07 | **0.82** | 0.14 |
| b) | NE | 0 | 3 | **0.23** | 0.40 | **0.45** | 0.39 | **0.34** | 0.34 | **0.43** | 0.42 | **0.45** | 0.36 | **0.61** | 0.27 | **0.77** | 0.16 |
|  |  | 1 | 6 | **0.26** | 0.37 | **0.43** | 0.41 | **0.50** | 0.45 | **0.54** | 0.44 | **0.61** | 0.35 | **0.73** | 0.17 | **0.74** | 0.18 |
| c) | GOL | 0 | 4 | **0.20** | 0.08 | **0.23** | 0.08 | **0.29** | 0.08 | **0.42** | 0.12 | **0.53** | 0.20 | **0.59** | 0.12 | **0.74** | 0.19 |
|  |  | 1 | 6 | **0.32** | 0.09 | **0.46** | 0.18 | **0.53** | 0.11 | **0.55** | 0.16 | **0.65** | 0.04 | **0.70** | 0.02 | **0.77** | 0.15 |
| d) | MIT | 0 | 3 | **0.76** | 0.24 | **0.86** | 0.08 | **0.87** | 0.07 | **0.89** | 0.09 | **0.96** | 0.04 | **0.97** | 0.03 | **0.94** | 0.07 |
|  |  | 1 | 4 | **0.51** | 0.35 | **0.56** | 0.32 | **0.68** | 0.33 | **0.73** | 0.31 | **0.94** | 0.05 | **0.96** | 0.04 | **0.97** | 0.04 |

## Supplementary Note 3: Hela-2

**Supplementary Fig. S7.**
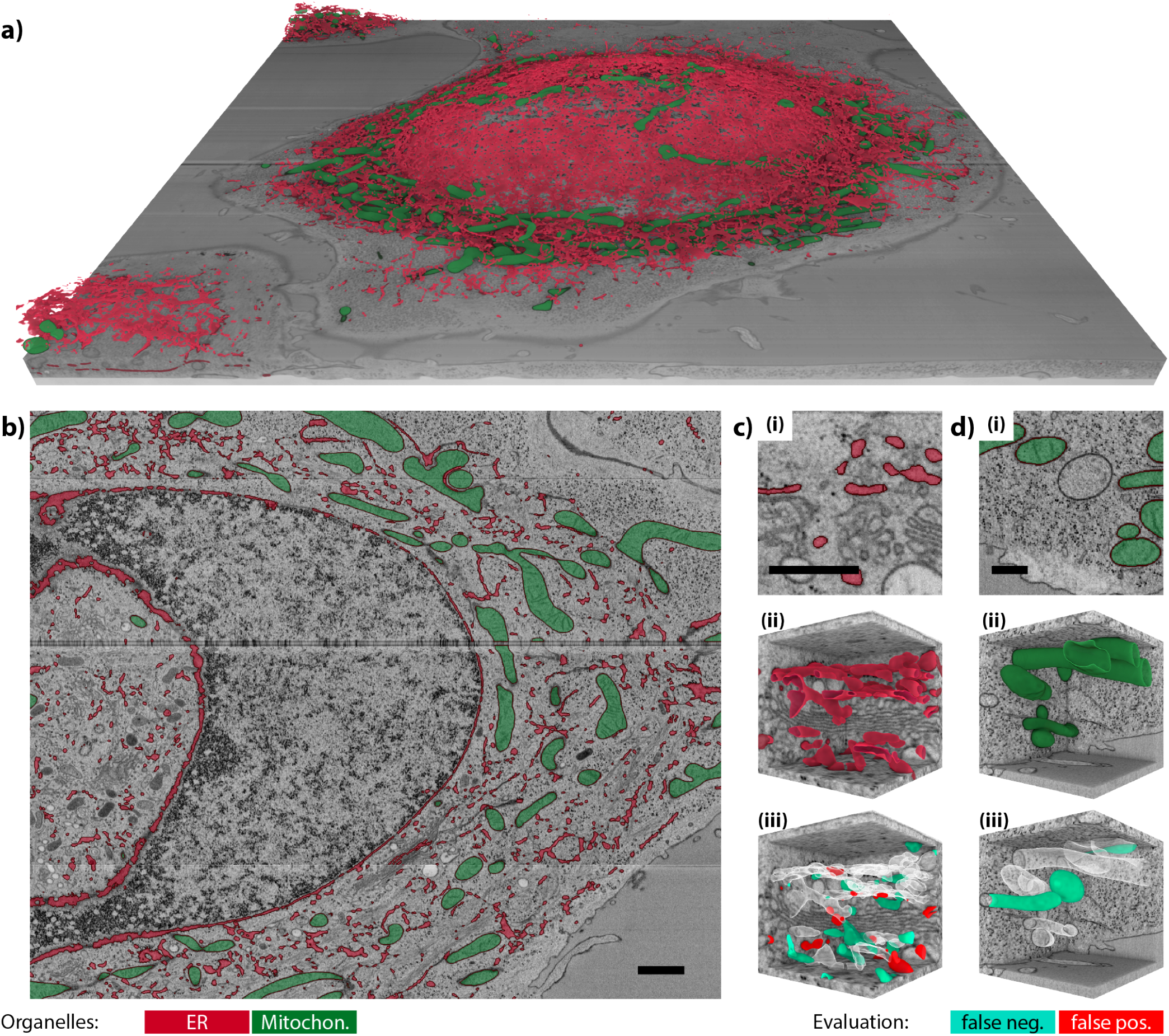
Segmentation results for Hela-2. The **Hela-2** dataset was annotated for ER (total of 12 subsets) and mitochondria (total of five subsets). In a) 3D visualization of the prediction result of the whole dataset. In b), a section of the dataset in the middle of the cell body, with the selected cell to a segment showing the predicted organelles with ER in red and mitochondria in green. In c) ER (red), the result of a validation cube where annotation was performed in native 4 nm isotropic resolution, and the final accuracy value of the cube (IoU) is 0.71. In d), mitochondria (green), annotated with a resolution of 10 nm isotropic, with an IoU of 0.75. IoU can vary among annotated cubes, from 0.75 up to 0.95, with the average value shown in Table S2.2. Scale bars are 2.5 µm (b) and 0.5 µm (c and d)

**Supplementary Table S2.1.**
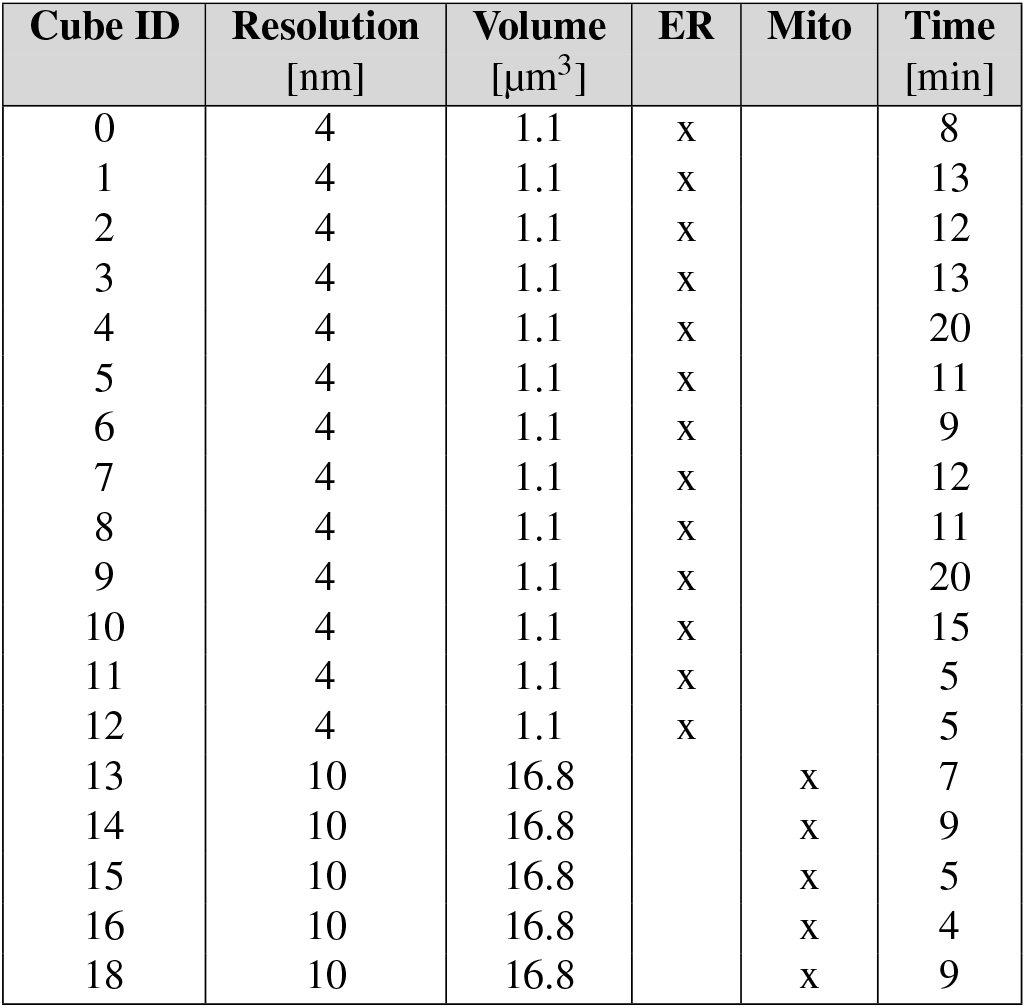
Annotated volumes and annotation time for each cube in Hela-2. Subset annotation of the **Hela-1** dataset for ER, NE, Golgi, and mitochondria (Mito). The columns refer to the cube ID, annotation resolution, cellular volume, the respective annotated organelles, and annotation time for each of the annotated subsets. The average time for annotating a cube was in the order of 14 min, depending mostly on the volume of organelle present on the cube and how many different classes were annotated on the same cube. The total volumes or annotated cubes were 23.1 µm^3^ for the ER, 12.6 µm^3^ for the NE and Golgi, each, and 67.2 µm^3^ for the mitochondria while the total cellular area was approximately 1800 µm^3^.

**Supplementary Table S2.2.** Evaluation of segmentation scores for the HeLa-2 dataset. In the table we show average values of IoU accuracy for a total of four validation cubes for ER and mitochondria, each. The columns show computations for different values of the parameter β ranging from 0.5 to 0.8. The highlighted columns correspond to the beta values used in the final segmentation shown in Figure S2. The maximum value of IoU might not always correspond to the chosen beta value, since the choice is also done based on visual inspection of the validation result and additionally takes the segmentation quality on the instance level into account which is not reflected in the IoU.

|  | beta: |  |  | 0.5 |  | 0.55 |  | 0.6 |  | 0.65 |  | 0.7 |  | 0.75 |  | 0.8 |  |
| --- | --- | --- | --- | --- | --- | --- | --- | --- | --- | --- | --- | --- | --- | --- | --- | --- | --- |
|  |  | It-ID | n_GT | mean | s.d. | mean | s.d. | mean | s.d. | mean | s.d. | mean | s.d. | mean | s.d. | mean | s.d. |
| **a)** | **ER** | 0 | 4 | **0.24** | 0.23 | **0.44** | 0.30 | **0.43** | 0.27 | **0.44** | 0.25 | **0.46** | 0.14 | **0.53** | 0.13 | **0.56** | 0.17 |
|  |  | 1 | 6 | **0.27** | 0.22 | **0.48** | 0.33 | **0.47** | 0.24 | **0.57** | 0.04 | **0.6** | 0.09 | **0.64** | 0.21 | **0.6** | 0.24 |
|  |  | 2 | 9 | **0.43** | 0.32 | **0.47** | 0.33 | **0.49** | 0.34 | **0.69** | 0.10 | **0.66** | 0.04 | **0.68** | 0.17 | **0.61** | 0.23 |
|  |  | 3 | 13 | **0.27** | 0.19 | **0.49** | 0.31 | **0.67** | 0.16 | **0.73** | 0.09 | **0.69** | 0.08 | **0.68** | 0.13 | **0.62** | 0.22 |
| **d)** | **MIT** | 0 | 4 | **0.71** | 0.2 | **0.84** | 0.13 | **0.84** | 0.13 | **0.86** | 0.13 | **0.85** | 0.14 | **0.88** | 0.11 | **0.88** | 0.11 |
|  |  | 1 | 5 | **0.74** | 0.18 | **0.84** | 0.13 | **0.84** | 0.13 | **0.85** | 0.14 | **0.87** | 0.13 | **0.88** | 0.11 | **0.89** | 0.12 |

## Supplementary Note 4: Calu-3

**Supplementary Fig. S8.**
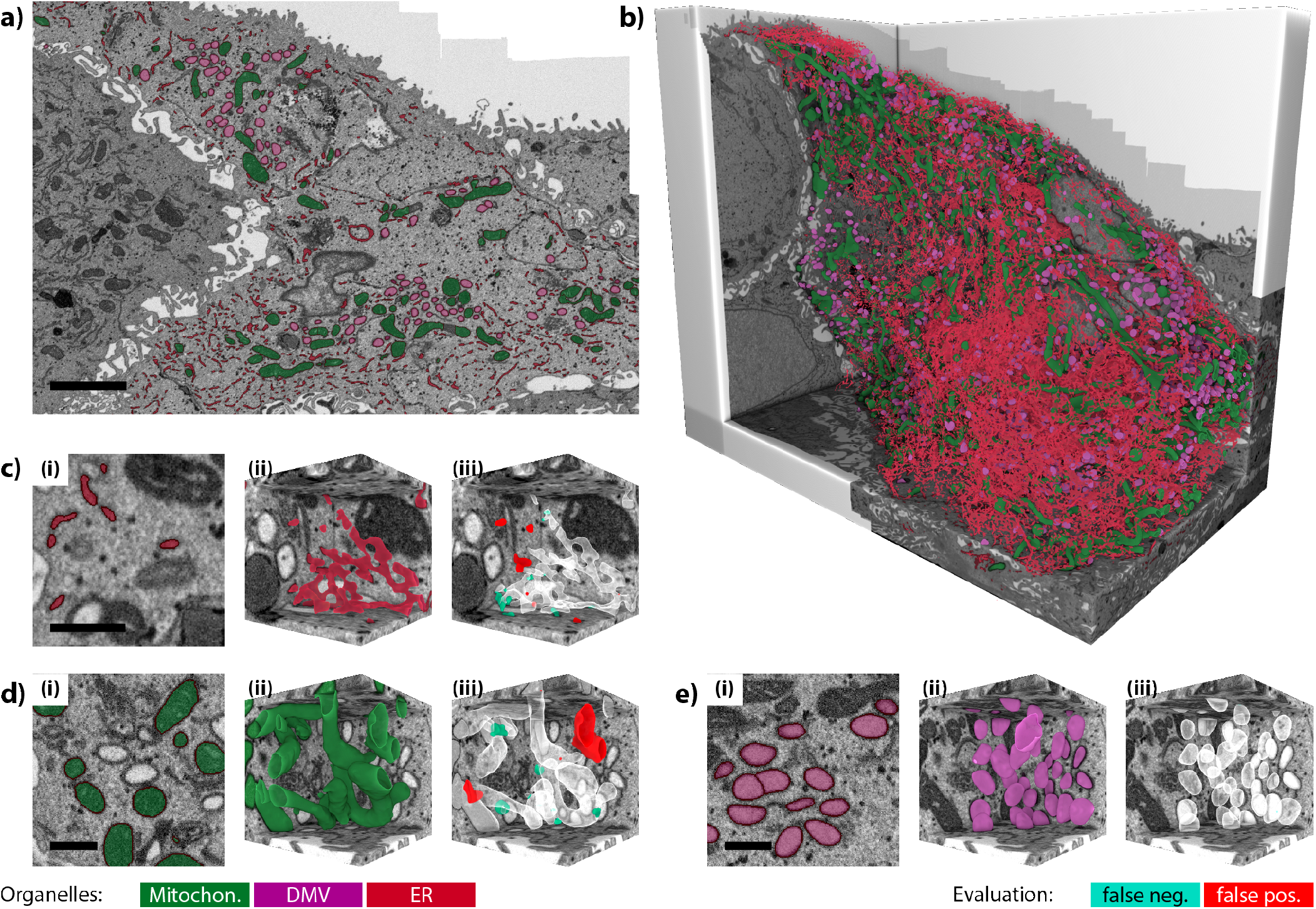
Segmentation results on Calu3-3. The **Calu3-3** dataset was annotated for ER (total of three subsets), DMVs (total of five subsets), and mitochondria (total of five subsets). In a), a section of the selected cell showing the predicted organelles with ER in red, mitochondria in green and DMVs in pink. The outer cells were masked out. In b) the full view of the resulting segmentation in 3D. c) to e) Results on validation cubes with (i) an example slice, (ii) the 3D representation of the segmentation, and (iii) the difference to the annotated ground truth: c) ER (red) annotation was performed in 5 nm isotropic resolution yielding an IoU of 0.88 in the shown cube. Mitochondria (green) and DMVs (purple) were annotated jointly in the native dataset resolution of 8 nm isotropic, with d) the annotation of the mitochondria and an IoU of 0.57 in the shown cube and e) the DMVs with an IoU of 0.99. Scale bars are 2.5 µm (a) and 0.5 µm (c to e)

**Supplementary Table S3.1.** Annotated volumes and annotation time for each cube in Calu3-3. The columns refer to the cube ID, annotation resolution, cellular volume, the respective annotated organelles, and annotation time for each of the annotated subsets. The average annotation time of the five DMV/Mitochondria subsets was around 18 minutes covering a total volume of 42.9 µm^3^ and the average annotation time for the three ER subsets was around 21 minutes covering a volume of 6.3 µm^3^. The full volume of the target cell covered by the dataset was approximately 3600 µm^3^.

| Cube ID | Resolution [nm] | Volume [$\mu\text{m}^3$] | ER | DMV | Mito | Time [min] |
| --- | --- | --- | --- | --- | --- | --- |
| 0 | 5 | 2.1 | x |  |  | 17 |
| 1 | 5 | 2.1 | x |  |  | 22 |
| 2 | 5 | 2.1 | x |  |  | 17 |
| 3 | 8 | 8.6 |  | x | x | 17 |
| 4 | 8 | 8.6 |  | x | x | 21 |
| 5 | 8 | 8.6 |  | x | x | 30 |
| 6 | 8 | 8.6 |  | x | x | 17 |
| 7 | 8 | 8.6 |  | x | x | 19 |

**Supplementary Table S3.2.** Evaluation of segmentation scores for the HeLa-2 dataset. In the table we show average values of IoU accuracy for a total of four validation cubes for ER and mitochondria, each. The columns show computations for different values of the parameter β ranging from 0.5 to 0.8. The highlighted columns correspond to the beta values used in the final segmentation shown in Figure S2. The maximum value of IoU might not always correspond to the chosen beta value, since the choice is also done based on visual inspection of the validation result and additionally takes the segmentation quality on the instance level into account which is not reflected in the IoU.

|  | beta: |  |  | 0.5 |  | 0.55 |  | 0.6 |  | 0.65 |  | 0.7 |  | 0.75 |  | 0.8 |  |
| --- | --- | --- | --- | --- | --- | --- | --- | --- | --- | --- | --- | --- | --- | --- | --- | --- | --- |
|  |  | It-ID | n_GT | mean | s.d. | mean | s.d. | mean | s.d. | mean | s.d. | mean | s.d. | mean | s.d. | mean | s.d. |
| a) | ER | 0 | 3 | **0.75** | 0.02 | **0.82** | 0.03 | **0.76** | 0.12 | **0.78** | 0.15 | **0.74** | 0.16 | **0.74** | 0.14 | **0.65** | 0.05 |
| b) | MIT | 0 | 3 | **0.59** | 0.42 | **0.57** | 0.40 | **0.59** | 0.40 | **0.68** | 0.25 | **0.68** | 0.25 | **0.68** | 0.23 | **0.66** | 0.16 |
|  |  | 1 | 5 | **0.75** | 0.17 | **0.77** | 0.18 | **0.78** | 0.20 | **0.81** | 0.13 | **0.80** | 0.14 | **0.79** | 0.13 | **0.78** | 0.13 |
| c) | DMV | 0 | 3 | **0.99** | 0.01 | **0.99** | 0.01 | **0.99** | 0.01 | **0.99** | 0.01 | **0.98** | 0.03 | **0.97** | 0.02 | **0.96** | 0.02 |
|  |  | 1 | 5 | **0.93** | 0.06 | **0.93** | 0.06 | **0.98** | 0.02 | **0.99** | 0.01 | **0.99** | 0.01 | **0.99** | 0.01 | **0.99** | 0.01 |

## Supplementary Note 5: Macrophage-4

**Supplementary Fig. S9.**
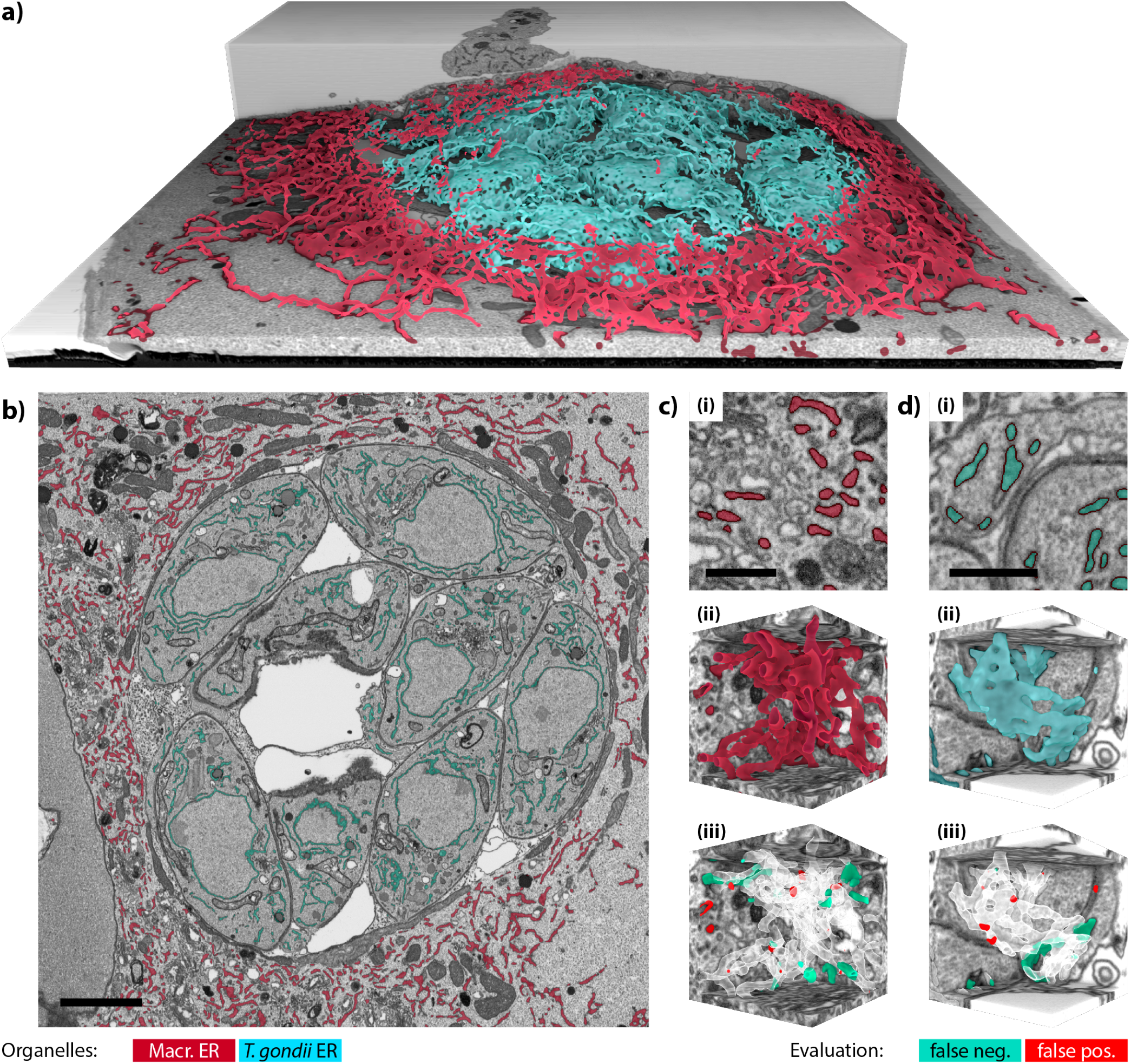
Segmentation results on Macrophage-4. The **Macrophage-4** dataset was annotated exclusively for ER, however individually for the ER of the macrophage (two subsets) and the ER of the *T. gondii* parasites (four subsets). a) The full 3D context of the macrophage ER (red) and the parasite ER (cyan). b) A top-view ortho-slice of the dataset showing the two ER types as an overlay. c) and d) examples of validation cubes used to determine the segmentation quality which was not part of the training set, with (i) an example slice with ER overlay, (ii) the ER in the cube in its 3D representation, and (iii) the difference to the annotation validation ground truth, for this cube with an IoU of 0.87 for cell ER and 0.86 for the toxoplasma parasite ER. c) Validation cube for the macrophage ER, d) validation cube for the *T. gondii* ER. Scale bars are 2.5 µm (b) and 0.5 µm (c and d)

**Supplementary Table S4.1.** Annotated volumes and annotation time for each cube in Macrophage-4 and *T. gondii* ER. The columns refer to the cube ID, annotation resolution, cellular volume, the respective annotated organelles, and annotation time for each of the annotated subsets. The average annotation time for the macrophage ER was 17 minutes covering 4.2 µm^3^ at the native dataset resolution of 5 nm isotropic. For the region of the *T. gondii* ER, the average annotation time was around 8.0 minutes covering a total volume of 4.3 µm^3^. We differentiated between ER in the outer cell and the *T. gondii* parasites due to the difference in ER morphology. The total volume of the macrophage including the parasites was 880 µm^3^.

| Cube ID | Resolution [nm] | Volume [$\mu\text{m}^3$] | ER mac | ER tox | Time [min] |
| --- | --- | --- | --- | --- | --- |
| 0 | 10 | 16.8 | x |  | 22 |
| 1 | 10 | 16.8 | x |  | 5 |
| 2 | 10 | 16.8 |  | x | 10 |
| 3 | 10 | 16.8 |  | x | 7 |
| 4 | 10 | 16.8 |  | x | 9 |
| 5 | 10 | 16.8 |  | x | 6 |

**Supplementary Table S4.2.** Evaluation of segmentation scores by means of IoU for the Macrophage-4 dataset, for ER, both cellular and *T. gondii* parasites. In the table, the average value of IoU accuracy, for a total of three validation cubes, and for a range of beta values from 0.5 to 0.8. The highlighted columns correspond to the beta values used in the final segmentation shown in Figure S3. The maximum value of IoU might not always correspond to the chosen beta value, since the choice is also done based on visual inspection of the validation result and additionally takes the segmentation quality on the instance level into account which is not reflected in the IoU.

|  | beta: |  |  | 0.5 |  | 0.55 |  | 0.6 |  | 0.65 |  | 0.7 |  | 0.75 |  | 0.8 |  |
| --- | --- | --- | --- | --- | --- | --- | --- | --- | --- | --- | --- | --- | --- | --- | --- | --- | --- |
|  |  | It-ID | n_GT | mean | s.d. | mean | s.d. | mean | s.d. | mean | s.d. | mean | s.d. | mean | s.d. | mean | s.d. |
| a) | MAC | 0 | 2 | **0.73** | 0.06 | **0.82** | 0.03 | **0.84** | 0.04 | **0.86** | 0.01 | **0.88** | 0.01 | **0.86** | 0.01 | **0.84** | 0.01 |
| b) | T.g. | 0 | 2 | **0.53** | 0.48 | **0.81** | 0.10 | **0.80** | 0.13 | **0.79** | 0.19 | **0.79** | 0.22 | **0.79** | 0.21 | **0.71** | 0.14 |
|  |  | 1 | 4 | **0.87** | 0.13 | **0.90** | 0.11 | **0.91** | 0.09 | **0.93** | 0.06 | **0.91** | 0.08 | **0.90** | 0.05 | **0.88** | 0.04 |

## Supplementary Note 6: Platynereis-5

**Supplementary Fig. S10.**
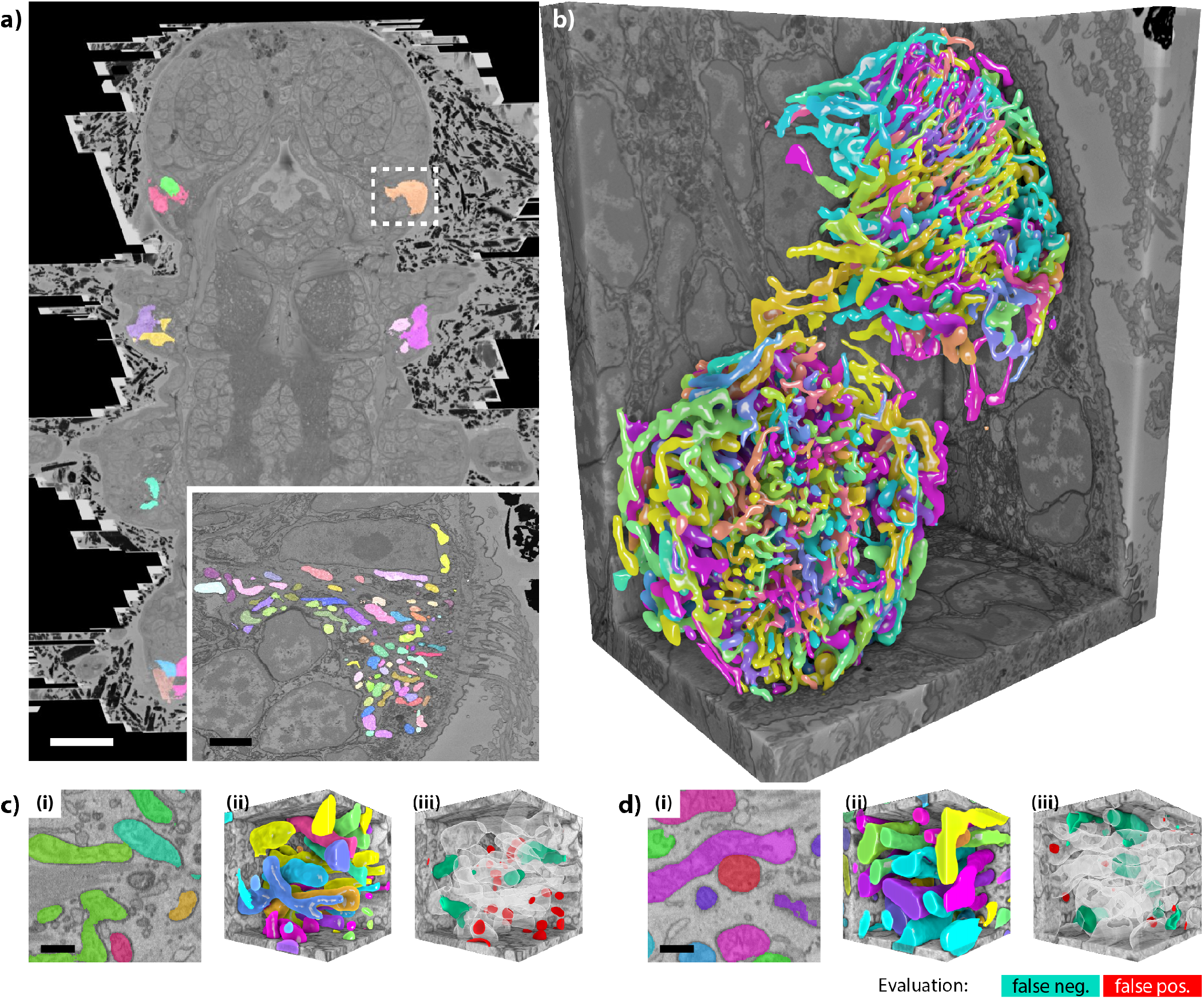
Segmentation results on Platynereis-5. The **Platynereis-5** dataset was annotated for mitochondria in ciliated cells using 10 nm isotropic resolution. The ciliated cells were selected using a cell segmentation and cell type assignment from previous work (Vergara et al., 2021). a) Overview over the entire animal with a set of segmented ciliated cells. The inset shows a slice of a mitochondria segmentation of the cell indicated by the dashed white box. b) 3D rendering of the mitochondria segmentation of the cell highlighted in panel (a) and its neighboring ciliated cell. c) and d) show two of four validation cubes with (i) an example slice, (ii) the volume representation of the segmentation and (iii) the difference to the annotated ground truth. The respective segmentation accuracies were an IoU of 0.88 and 0.79, respectively. Scale bars are 25 µm (a), 2.5 µm (a, inset) and 0.5 µm (c and d)

**Supplementary Table S5.1.** Annotated volumes and annotation time for each cube for mitochondria in ciliated cells. The columns refer to the cube ID, annotation resolution, cellular volume, the cell ID in the cell segmentation, and annotation time for each of the annotated subsets. Each annotated cube had a length of 2.6 um, so each cube had a volume of 2.56 × 2.56 × 2.56 = 16.8 µm^3^. We annotated seven cubes which took around 22 minutes each, covering a total volume of 117 µm^3^. The full volume of the 31 segmented ciliated cells was 41000 µm^3^.

| Cube ID | Resolution<br>[nm] | Volume<br>[$\mu\text{m}^3$] | Mito | Time<br>[min] |
| --- | --- | --- | --- | --- |
| 0 | 10 | 16.8 | x | 28 |
| 1 | 10 | 16.8 | x | 20 |
| 2 | 10 | 16.8 | x | 33 |
| 3 | 10 | 16.8 | x | 8 |
| 4 | 10 | 16.8 | x | 26 |
| 5 | 10 | 16.8 | x | 23 |

**Supplementary Table S5.2.** Evaluation of segmentation scores by means of IoU for the Platynereis-5 dataset, for mitochondria. In the table, the average value of IoU accuracy, for a total of three validation cubes, and for a range of beta values from 0.5 to 0.8. The highlighted columns correspond to the beta values used in the final segmentation shown in Figure S3. The maximum value of IoU might not always correspond to the chosen beta value, since the choice was also based on visual inspection of the validation result and additionally takes the segmentation quality on the instance level into account which is not reflected in the IoU. A value of β = 0.5 performed best in terms of instance segmentation while yielding an IoU close to the maximum achieved by slightly higher values of β.

| beta: |  | 0.5 |  | 0.55 |  | 0.6 |  | 0.65 |  | 0.7 |  | 0.75 |  | 0.8 |  |
| --- | --- | --- | --- | --- | --- | --- | --- | --- | --- | --- | --- | --- | --- | --- | --- |
| It-ID | n_GT | mean | s.d. | mean | s.d. | mean | s.d. | mean | s.d. | mean | s.d. | mean | s.d. | mean | s.d. |
| 0 | 5 | **0.67** | 0.39 | **0.74** | 0.36 | **0.78** | 0.30 | **0.82** | 0.21 | **0.85** | 0.17 | **0.87** | 0.08 | **0.83** | 0.14 |
| 1 | 7 | **0.87** | 0.06 | **0.89** | 0.04 | **0.89** | 0.03 | **0.82** | 0.14 | **0.85** | 0.17 | **0.88** | 0.17 | **0.86** | 0.17 |

## Supplementary Note 7: Computation of supervoxels

**Supplementary Fig. S11.**
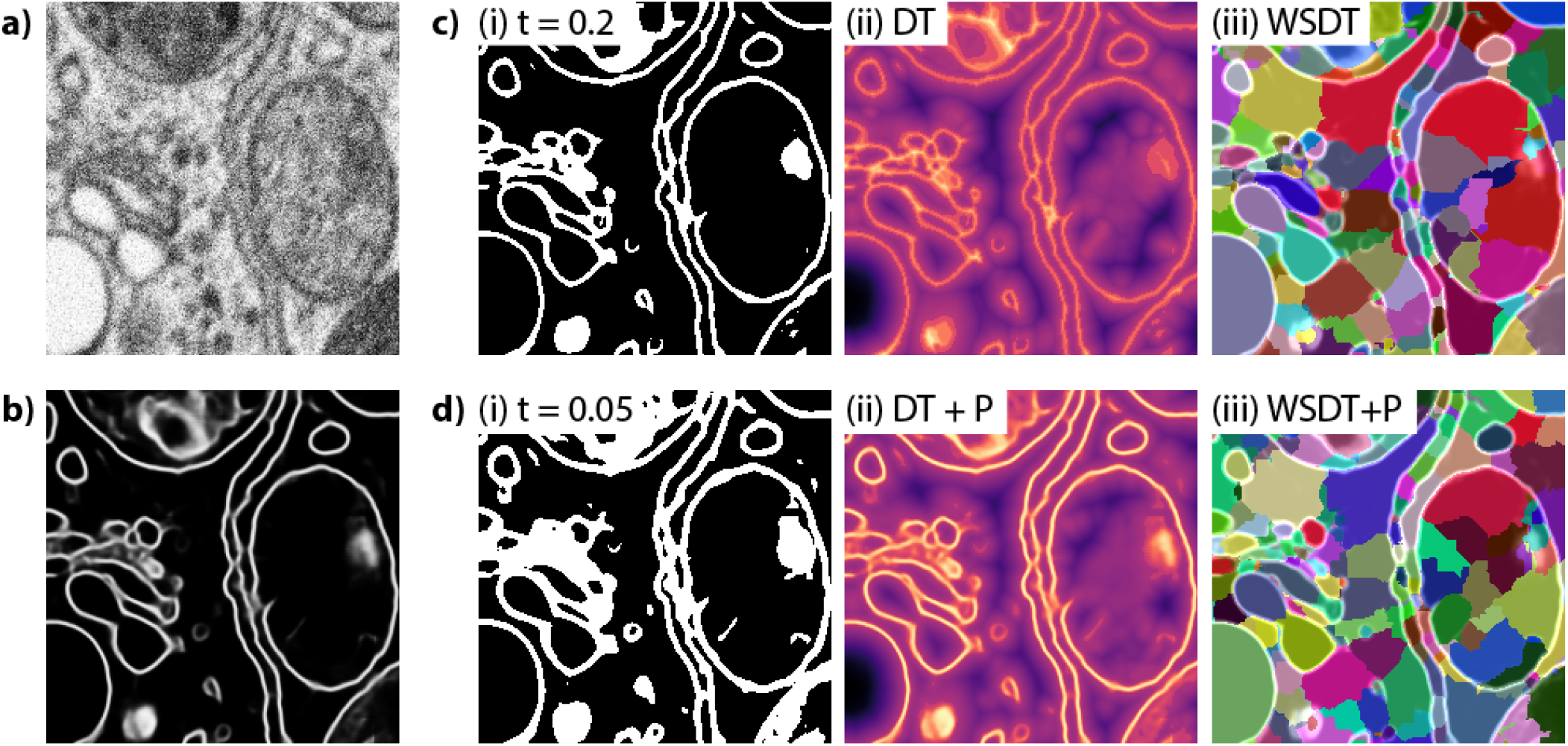
Computation of supervoxels from the membrane probability map. a) Raw input data used to compute b) the membrane probabilities. c) The workflow of supervoxel preparation proposed by Beier et al. 2017, where the data is thresholded (i), a signed boundary distance transform is computed (ii), and supervoxels are generated by a watershed based on the distance transform (iii). d) The workflow of the supervoxel computation as used within the CebraEM workflow consists of (i) masking the membrane probabilities with a low threshold, (ii) generation of a composite map of the probabilities, and a distance transform computed on the mask which yields the basis for the seeded watershed computation producing (iii) the final supervoxels.

## Supplementary Note 8: CebraANN graphical user interface

**Supplementary Fig. S12.**
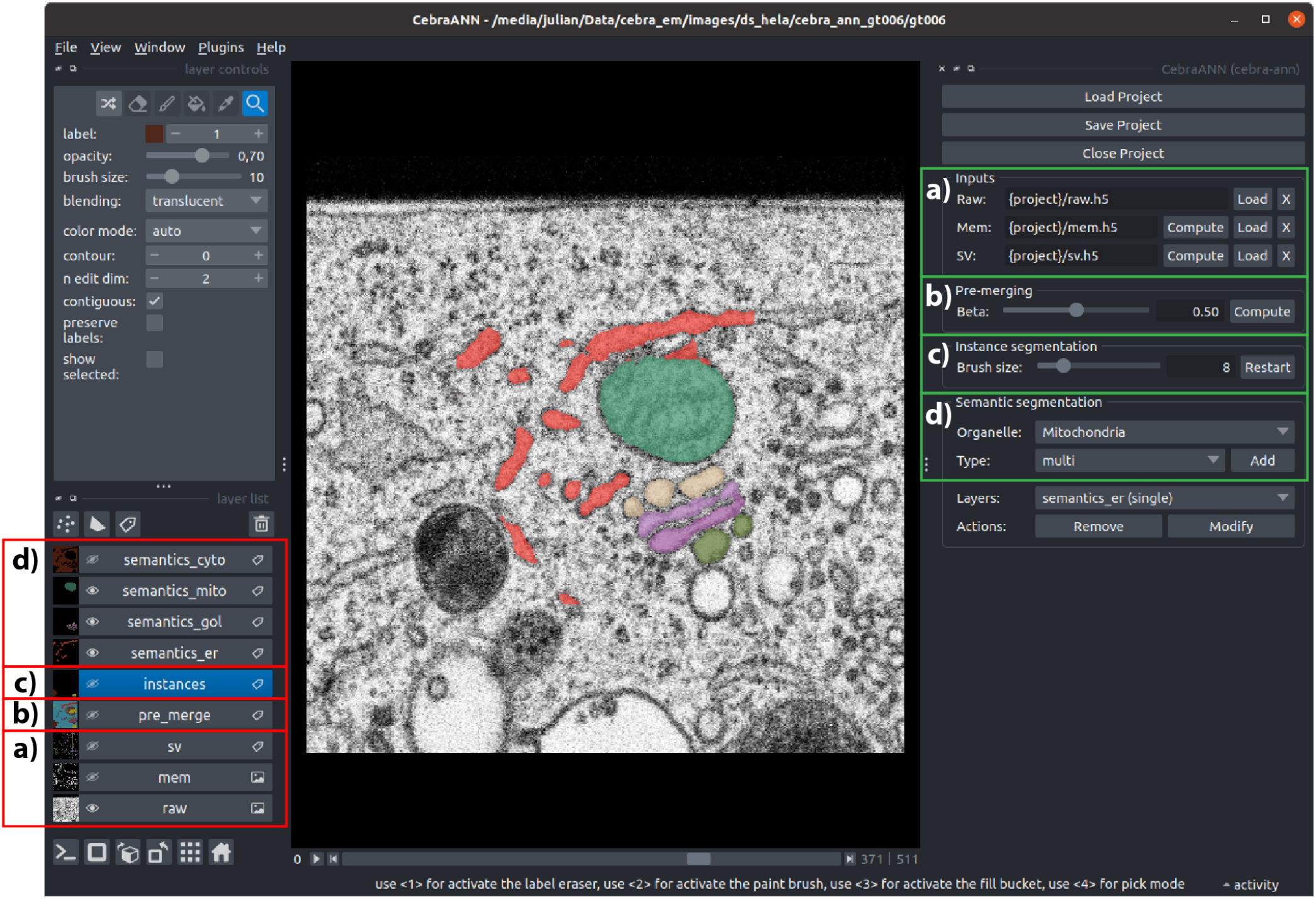
Graphical user interface of the CebraANN Napari plugin. Green frames indicate the workflow settings of the different steps: a) Input loading (Raw as raw data, Mem as membranes and SV as supervoxels computed in the previous step of the workflow), b) pre-merging to select the beta value and decide the initial partition state of the supervoxels, c) instance segmentation, the size of the brush for correcting a selected instance value, and d) semantic segmentation, where the class of organelle to segment is selected. The red frames show the respective data layers corresponding to each workflow step (a to d, respectively), so the user can select them for visualization purposes.

## Supplementary Note 9: ER collapse affects the segmentation

**Supplementary Fig. S13.**
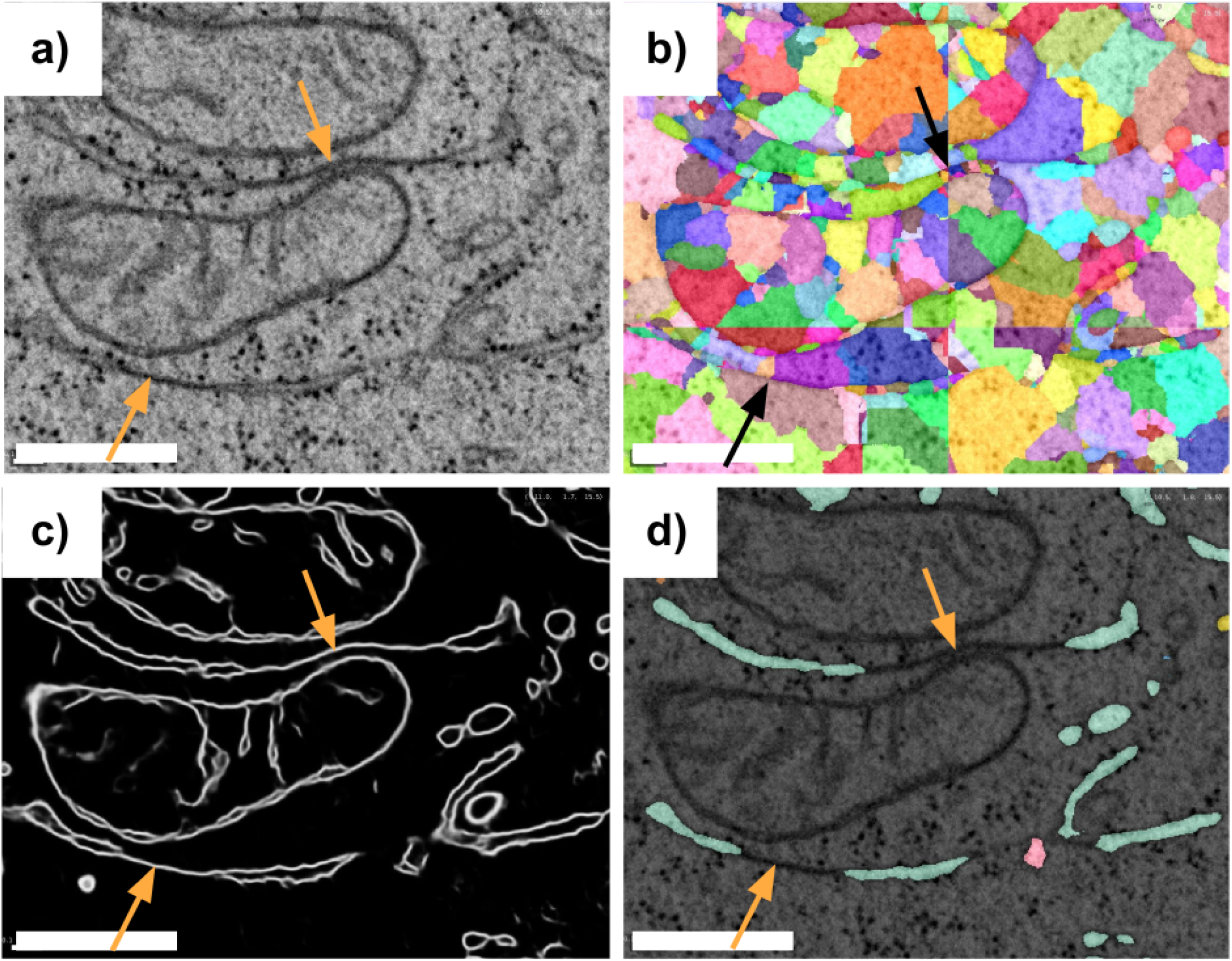
Illustration of ER collapse and the impact on the segmentation. a) Raw data showing instances of collapsed ER (arrows), b) where the membrane prediction recovers only one stretch of the membrane such that c) no supervoxels can be generated and d) the ER is not segmented here. Scale bars: 0.5 µm

